# Ancient genomes chronicle the history of adaptive evolution in East Asia

**DOI:** 10.64898/2026.10.16.759767

**Authors:** Dan Ju, Masixuan Zhao, Chuyan Li, Xinyu Wei, Mark Stoneking, Tianxiang Liu, Xiaotian Feng, Peng Cao, Qiaomei Fu

## Abstract

The transition from hunting and gathering to farming brought about novel selection pressures related to metabolism and immunity, but for East Asian populations the evolutionary genetic dynamics of these adaptations and their relationships to agriculture remain unclear. Using one of the largest datasets of ancient East Asian genomes (N=2236), we performed a genome-wide scan to identify selected variants and reconstructed their allele frequency changes over time and space. We tested for genetic association with farming using an environmental variable to represent historical farming intensity based on a large database of archaeobotanical sites (N=1220). Our analyses indicate selection began prior to the Neolithic for several well-established adaptive loci such as *FADS1*, *FUT2*, and *OCA2*. For rs1229984 at *ADH1B*, the strongest selection signal in our genome-wide scan, we observed variation in timing and strength of selection in East Eurasia and found evidence for a southern origin. Furthermore, *ADH1B*48His* was positively associated with wheat farming. Given these results, we propose a novel hypothesis about the fitness advantage of this variant. Our work underscores the heterogeneity in the strength and timing of selection across East Eurasia and provides a quantitative framework for testing agriculture’s role as a selection pressure.

## Introduction

Ancient DNA (aDNA) has greatly informed our understanding of human adaptation in regard to the traits influencing fitness and their corresponding selection pressures by providing snapshots of genotypes across time and space. As most of the aDNA has been sampled from Holocene Eurasia, the loci showing genomic signals of selection have often been linked to a suite of novel selection pressures associated with the transition from hunting and gathering to farming. Increased population density and greater proximity to domesticated animals increased exposure to pathogens; changes in diet led to some nutritional deficits while potentially providing an abundance of calories that could be unstable depending on local conditions^1^.

Previous studies have used ancient genomes to investigate selection primarily in West Eurasia^2–4^, but less is known about East Asia. The evolutionary histories of major adaptive loci in East Asia remain unclear due to the historically limited number of samples from East Asia, particularly from the south. While previous work has indicated broad sharing of selective sweeps in the last ten thousand years between West and East Eurasia^5,6^, it is unknown how uniform the timing and intensity of these sweeps were within East Eurasia itself. East Eurasia is at least twice as large as West Eurasia and varies greatly in environmental factors of relevance to fitness, such as climate and arable land. Claims about agriculture-driven selection have been largely based on the timing of selection coinciding with major transitions related to subsistence in the Holocene. A limitation of this kind of evidence, however, is that the adoption of farming by various populations in Eurasia was gradual and uneven in space and time. In the case of the East Asian Neolithic, which began ∼10,000 years before present (BP), cereal crops of independent provenance—rice (*Oryza sativa*) along the Yangtze River Basin as well as broomcorn millet (*Panicum miliaceum*) and foxtail millet (*Setaria italica*) in the north—spread along different spatiotemporal trajectories, generating a dynamic landscape of crop cultivation across East Asia that profoundly changed in some areas with the later arrival of western domesticates such as wheat (*Triticum* spp.) and barley (*Hordeum vulgare*). This shifting agricultural landscape provides a natural experiment for testing the role of agriculture as a selective pressure.

Taking advantage of the recent growth of ancient genomes sequenced from East Asia, we performed the largest genome-wide scan of positive selection with data from this region to date. We provide insights into the timing and strength of selection and explore the spatial dynamics of adaptive variation. Furthermore, we tested hypotheses about agriculture driving selection on selected variants by applying a new approach that leverages the variation in agricultural intensification across time and space on datasets of ancient genomes (N=2236) and archaeobotanical site observations primarily from East Asia (N=1220). These datasets provide the basis for a comprehensive investigation into East Asian adaptive history.

## Results

### Preparation and imputation of ancient genomes from East Eurasia

We assembled a database of ancient nuclear genomes from East Eurasia from published studies (N=2000) and newly sequenced genomes sampled from North-Central China (N=85) and the lower Yangtze River (N=151), which we merged with present-day HGDP individuals from East Eurasia (N=233), to reconstruct allele frequency changes over time. Spatially, these data represent a large geographical transect covering Western Siberia to the Japanese archipelago to Southeast Asia (Fig. 1A). These ancient data ranged in time from 40,000 BP to 139 BP with the vast majority of individuals having lived in the past ten thousand years (Fig. 1B). The majority of the ancient genomes underwent enrichment by 1240k capture panel or Twist panel (N=1667) and around a quarter underwent shotgun-sequencing without enrichment (N=565). PCA with ancient genomes projected onto present-day genetic variation captured the major axes of ancestry in East Eurasia, including those representative of Yellow River, Northeast Asian, and Southern East Asian populations (Fig. 1C).

**Figure 1.**
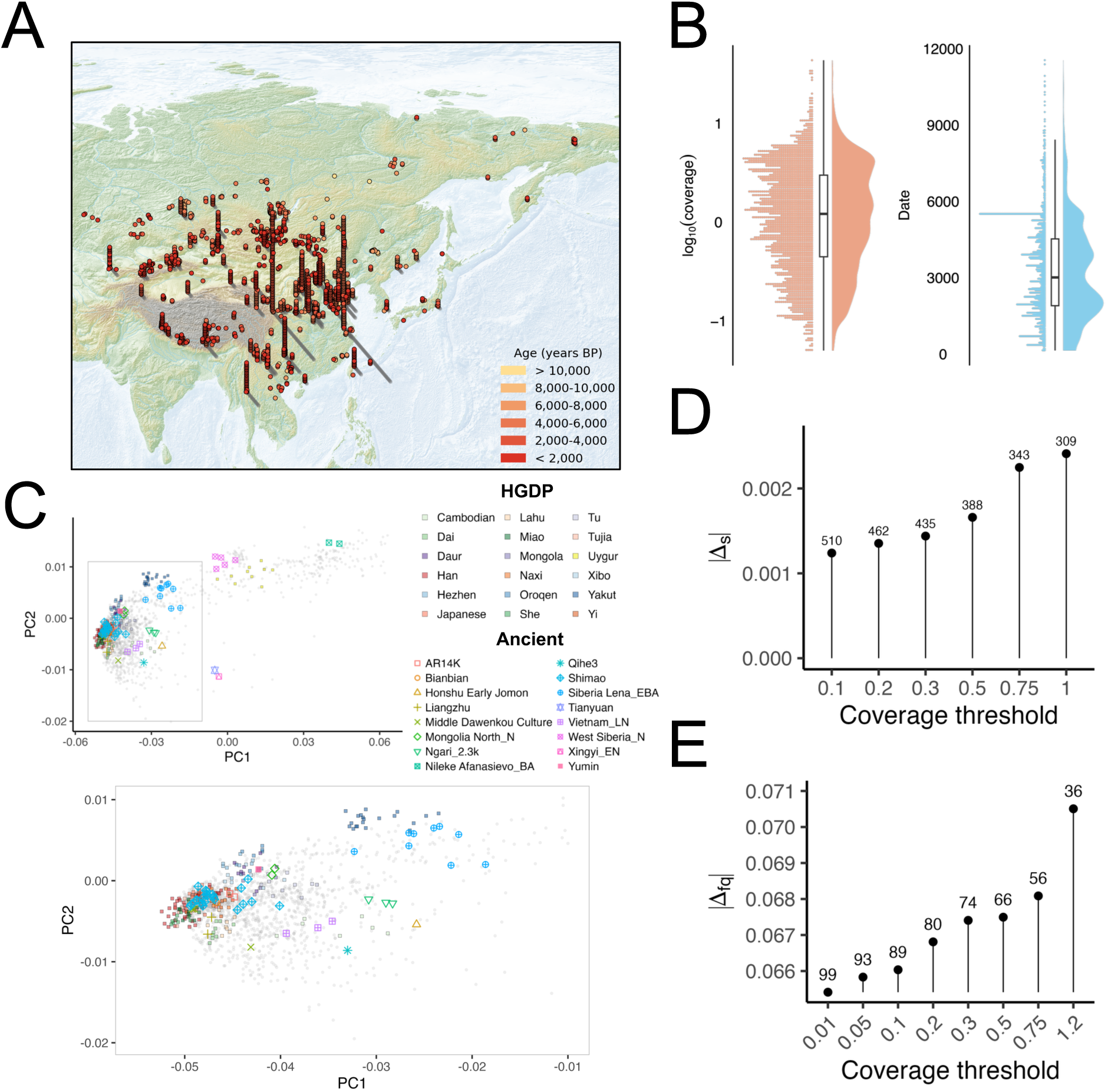
**Overview of the ancient East Asian dataset and imputation metrics**. (A) Locations of ancient individuals colored by time period. Individuals from the same location are stacked on top of each other. (B) Distributions of the radiocarbon dates of the samples (cut off at start of Holocene) and their genomic coverages at 1240k sites. (C) Genomic PCA based on imputed genotypes from GLIMPSE2 of ancient and present-day samples in the dataset. Select ancient individuals are highlighted and modern HGDP individuals are represented by filled boxes. Imputation accuracy evaluated by simulations based on median differences in (D) estimated selection coefficients between validation dataset and subsets of data based on different coverage thresholds for downsampled ancient capture genomes and (E) allele frequencies between high coverage HGDP Yakut genomes and subsets of downsampled genomes filtered by various coverage thresholds.

To increase the power of our dataset for downstream selection analyses, we performed genomic imputation with GLIMPSE2 using high-coverage data from globally diverse participants in the 1000 Genomes Project. We investigated the performance of imputation for ancient individuals through downsampling experiments (see Methods) to determine a reasonable coverage threshold for our selection analyses. In general, we observed decent genotype accuracy (r^2^ > 0.8) for imputation of common variants in individual East Eurasian capture-enriched genomes of at least 1x coverage (see Supplementary Note). Genotype accuracy, however, declined steeply even for common variants with decreasing coverage. Given that our analyses for studying selection are group-based, we considered how coverage thresholds influenced the accuracy of estimates of allele frequency and selection coefficients. Through two separate simulations that were tailored to the coverage distributions in our datasets, we found that using relatively permissive thresholds for our data led to slightly more accurate estimates when calculating population allele frequencies from shotgun data (Fig. 1D) and rates of allele frequency change estimated from time series of higher coverage ancient samples that were capture-enriched (Fig. 1E). Thus, for downstream analyses we mainly used a coverage threshold of >0.1x and >0.05x for capture and shotgun samples, respectively.

### Time series selection scan reveals adaptive variants

Several previous studies of selection using ancient East Asian genomes^4,6–8^ have relied on small sample sizes (N<300) and mostly been confined to SNPs on the 1240K array. One recent study^5^ has greatly increased the power of previous selection scans with a much larger sample of imputed ancient individuals from East Eurasia (N = 1812), but a substantial number appear to have originated outside of East Asia proper. To address the gap in sample size of ancient genomes from East Asia, we tested 7,135,256 SNPs in our dataset of imputed ancient and present-day genomes primarily from East Asia (N=2469) for unexpectedly large changes in frequency over time with a logistic regression model that controlled for potential confounders, including ancestry. We used a conventional genome-wide threshold of 5x10^-8^ to declare significance for SNP P-values that were adjusted with genomic control given test statistic inflation (λ = 2.57) (Supplementary Fig. 1). Our scan for positive selection yielded eight independent loci that were genome-wide significant (Fig. 2A). These outliers include established candidates for selection—the *ADH* gene cluster, the *HLA* region, a locus encompassing *ALDH2*, and *OCA2* (Supplementary Fig. 2A-C)—and other candidate loci that have been suggested to be under positive selection in East Asians but lacked genome-wide significant evidence based on aDNA: *OAS1* and *FUT2* (Supplementary Fig. 2D,E). Signals that have not been highlighted previously in other studies are a missense variant in *RSPH4A* and an intronic variant in *STAG1* (Supplementary Fig. 2F,G). Our strict significance threshold missed well-established loci such as *FADS1/2* (Supplementary Fig. 2I), so we also considered the top 0.1 percentile of genomic windows as potential candidates (Supplementary Fig. 3). We then leveraged the spatiotemporal context of our ancient individuals to address longstanding questions about the timing and nature of adaptive evolution in East Asia.

**Figure 2.**
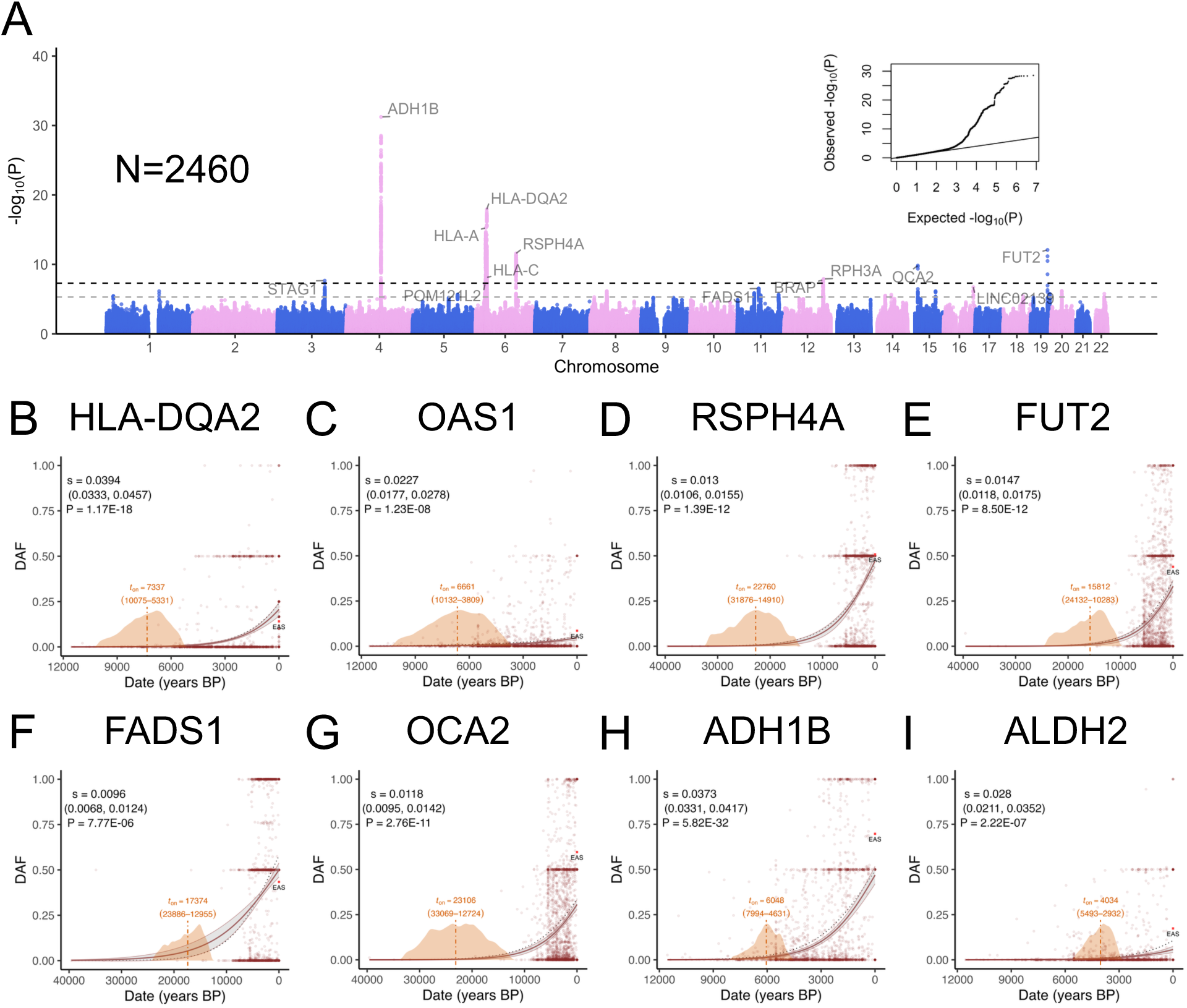
**Time series selection scan and allele frequency trajectories**. (A) Manhattan plot for the GC-corrected P-values for imputed SNPs of the date term from logistic regression of individuals from the last 12,000 BP. Dashed lines correspond to P-value thresholds of 5x10^-8^ and 5x10^-6^. Labelled genes were chosen based on the closest gene to the lowest P-value SNP position. (B-I) Predicted derived allele frequency (DAF) changes over time. The filled line with 95% CI envelope represents the regression with covariates including genomic PCs; the dashed line represents the prediction from a model with just date as predictor. On the X-axis the posterior distribution of the inferred changepoints is plotted with the median marked by the dashed vertical line and associated 95% credible interval. Dots represents the genotype dosages of each individual scaled down by half. At 0 BP, the allele frequency in 1000 Genomes East Asians, which were not included in the regression model, is plotted.

### Selection on immunity-related loci

Infectious disease has been hypothesized to have been an especially strong selective pressure during the Neolithic^9^. We observed a signal of selection at a key cluster of genes for the adaptive immune system, the human leukocyte antigen (HLA) region, which has been reported to be under positive selection and balancing selection in populations across the world^10^. rs9258176 near *HLA-DQA2* showed the most significant selection coefficient in the HLA region (s = 0.039; P = 1.17x10^-18^). Selection acted on the derived allele during the Neolithic starting from 7337 BP (95% CrI: 10,075 – 5331 BP) (Fig. 2B). Similarly, the derived allele of rs79230913 near *OAS1*, which encodes a protein important for cellular immunity against viral infection, showed evidence of selection during the Neolithic (s = 0.023; P = 1.23x10^-8^), beginning 6661 BP (95% CrI: 10,132 – 3809 BP) (Fig. 2C). This allele reached a frequency of 8.5 percent in present-day East Asians, whereas it is very rare to absent in other populations. *OAS1* has been suggested to be under independent episodes of positive selection in East and West Eurasian populations^11^, but previous selection scans of ancient East Eurasian genomes have not detected this signal. We found an older selective event at the missense variant rs784133 (Leu>His) in *RSPH4A* that began rising in frequency during the Upper Paleolithic (95% CrI: 31,876 – 14,910 BP) (Fig. 2D). This locus also appears to be an outlier in previous selection scans of ancient West and East Eurasians^2,5^. Selection on *RSPH4A*, which is involved in the structure and motility of cilia^12^, could be related to resistance against airborne infectious disease, as *RSPH4A* mutations increase infection risk by impairing the clearing of mucus in the lungs^13^, or to changes in fertility through oviduct cilia^14^. We observed other loci overlapping immune-related genes with evidence of selection, albeit not at the level of genome-wide significance, based on outlier genomic windows at the top 0.1 percentile of the empirical distribution: *FCAR*, *TNFRSF13C*, *TMEM229B*, *MUC2*/*MUC5AC*, *SYK*, and *GATA3* (Supplementary Fig. 6A-F). These loci generally appear to be under selection prior to the Holocene (Supplementary Fig. 7A-F).

While specific pathogen candidates driving selection at the aforementioned loci are unknown, nonsynonymous variants of the *FUT2* gene that cause the non-secretor phenotype have been demonstrated to protect against gastroenteritis-causing norovirus and rotavirus^15^. We observed selection for the missense allele at rs1047781 that is almost exclusively found in East Asian populations (s = 0.014; P = 2.44x10^-1^^1^). The onset of this selective event, which we estimated to be 15,812 BP (95% CrI: 24,132 – 10,283 BP), precedes the beginning of farming in East Asia (Fig. 2E). Pre-Neolithic selection was corroborated by CLUES2 using a two-epoch model (see Methods), which estimated s to be 0.035 (95% CrI: 0.031, 0.040) in the pre-Neolithic phase (Supplementary Fig. 8A). Our estimate of the timing of selection for the East Asian *FUT2* missense allele is considerably older than that of a previous study^15^, which observed that this allele only increased in frequency within the last 2000 years in East Asia. Interestingly, we also observed an increase in frequency for the West Eurasian truncation *FUT2* allele in the northwestern part of our dataset, which has an almost opposite spatial distribution compared to the East Asian variant (Supplementary Fig. 10 and Fig. 11). The onset of the truncation allele’s rise (95% CrI: 10,296, 8234 BP) overlaps with an estimate of ∼8700 BP reported in European populations^15^ (Supplementary Fig. 12). The difference in timing of selection between West and East Eurasia suggests either that the common selective pressure, such as gastroenteritis-causing pathogens, was prevalent earlier in East Asia or that a different selection pressure altogether existed in East Asia at the time.

### Pre-Neolithic rise of the derived FADS1/2 haplotype

The derived *FADS1/2* haplotype, associated with higher expression of these genes, is thought to have been evolutionarily advantageous for diets enriched in plants given their lower long-chain polyunsaturated fatty acids (LC-PUFAs) content, so this locus has been hypothesized to have evolved in response to agriculture. We observed evidence of selection at the *FADS* locus in our genome-wide scan, with the top SNP being rs174546 (s = 0.01; P = 7.77x10^-6^), the same SNP used to proxy the derived *FADS1/2* haplotype in other studies^16^. Previous reports based on East Asians have provided discordant timeframes for selection at *FADS*, with estimates ranging from very ancient (158,000 – 27,000 BP)^16^ to as recent as ∼6000 BP^5^. Our analysis indicates selection for the derived *FADS* haplotype began prior to the Neolithic, starting from 17,374 BP (95% CrI: 23,886 – 12,955 BP) (Fig. 2F). The inferred allele frequency trajectory from CLUES2 also showed an increase in the frequency before the Neolithic (Supplementary Fig. 9B), although the two-epoch model did not detect a significant selection coefficient in the pre-Neolithic phase (Supplementary Fig. 8B). This earlier timing of selection on *FADS*, while surprising based on the agricultural hypothesis, can still be attributed to dietary pressures in the absence of agriculture. Upper Paleolithic hunter-gatherers increasingly consumed flora such as tubers and Panicae grasses at the end of the last glacial maximum (LGM) in Northern China^17^. It has also been shown that Neanderthals carried the derived *FADS* haplotype^16,18^ and that the derived haplotype was anciently selected in Africa^19^.

### Evolution of the light skin pigmentation allele of OCA2

We detected a strong signal of selection for the derived light allele at the missense variant rs1800414 in *OCA2* (s = 0.012; P = 2.76 x10^-11^). While this variant was known to be highly differentiated in present-day East Asians compared to other populations^20^, previous selection scans using ancient East Eurasian genomes did not detect it, possibly due to inadequate representation of East Asian genomes. The selective sweep for the light allele at this locus began sometime in the Upper Paleolithic (95% CrI: 33,069 – 12,724 BP) (Fig. 2G). In contrast to the claim that most of the adaptive lightening of skin pigmentation occurred before the Holocene for East Asians^5^, the selection signal we detected at rs1800414 indicates that selection for lighter pigmentation persisted throughout the Neolithic. Consistent with the hypothesis that UVB radiation drove selection on skin pigmentation, we observed a positive relationship between the frequency of the light allele and latitude (P = 5.43 x 10^-3^), but the frequency of the light allele was actually more strongly associated with longitude (P = 9.32 x 10^-7^).

### Spatiotemporal dynamics of alcohol metabolism loci

Altered alcohol metabolism in East Asian populations can be attributed to the Arg48His variant (rs1229984) of *ADH1B*, which accelerates the oxidation of ethanol into acetaldehyde, and the Glu504Lys variant (rs671) of *ALDH2* that inhibits the breakdown of acetaldehyde, causing the red flush syndrome common in East Asians. The *ADH1B* locus exhibited the strongest signal in our selection scan (s = 0.037; P = 5.82x10^-32^). The adaptive haplotype has been estimated to have appeared roughly at the beginning of the Neolithic, and previous time series analyses suggest a selective sweep within the last ten thousand years^6,21^ with one study finding a sudden rise of the 48His allele around 4000-3000 BP^5^. In contrast, we estimated the onset of selection at 6048 BP (95% CrI: 7994 – 4631 BP), which, while later than the earliest evidence of rice domestication (∼11,000 BP)^22^ and alcohol production (∼10,000 BP)^23^ in China, places the start of selection in the middle rather than late Neolithic (Fig. 2H). Selection also acted on rs671 at *ALDH2* (s = 0.028; P = 2.22x10^-7^), although around 2000 years later (95% CrI: 5493 – 2932 BP) (Fig. 2I). More recent selection on rs671 compared to rs1229984 has been observed in other studies as well^7,21^.

Given the variation in timelines of agricultural transition across East Eurasia^24^, we investigated whether there were differences in the strength and timing of selection with respect to geography (Fig. 3A) (Supplementary Fig. 14). We observed spatial heterogeneity in selection coefficients for rs1229984 based on a linear mixed model testing for an interaction between region and time (P = 1.66x10^-4^). The 48His allele showed the most rapid change in the Loess Plateau & Ordos Plateau (LPOP) region where s = 0.078 (95% CI: 0.051, 0.104), which was significantly higher than that of the Huang He Plain (HHP) (P = 0.01) and Chang Jiang & Southern Coastal China (CJSC) (P = 0.003) regions (Fig. 3B). On the other hand, we observed no evidence of selection in the Qinghai-Tibetan Plateau (QTP) (P = 0.308), which is consistent with a previous study of present-day Tibetans^25^, and the northwestern regions of Junggar & Alashan Semi-deserts & Altai Mountains (JADAM) (P = 0.45) and Tianshan & Taklimakan Desert (TSTD) (P = 0.42). We found the earliest rise in derived allele frequency in the regions of Northeast Asia (NEA) followed by CJSC (Fig. 3C). The early timing in CJSC is consistent with the hypothesis that rice-farming populations around the Yangtze River were the source of the adaptive *ADH1B* haplotype in East Asians^26^. While we observed the earliest onset in NEA, we note that the regression model adjusting for ancestry suggests that this early rise could be explained by gene flow (Supplementary Fig. 17H). Nonetheless, our stratified analysis indicates that selection for the 48His allele likely occurred even earlier than indicated by our cohort-wide estimate, as selection began 8283 BP (95% CrI: 11,099-5978 BP) in the CJSC region (Supplementary Fig. 17B), closer in time to the earliest evidence of alcohol production at the early Neolithic site Shangshan in the lower Yangtze^23^.

**Figure 3.**
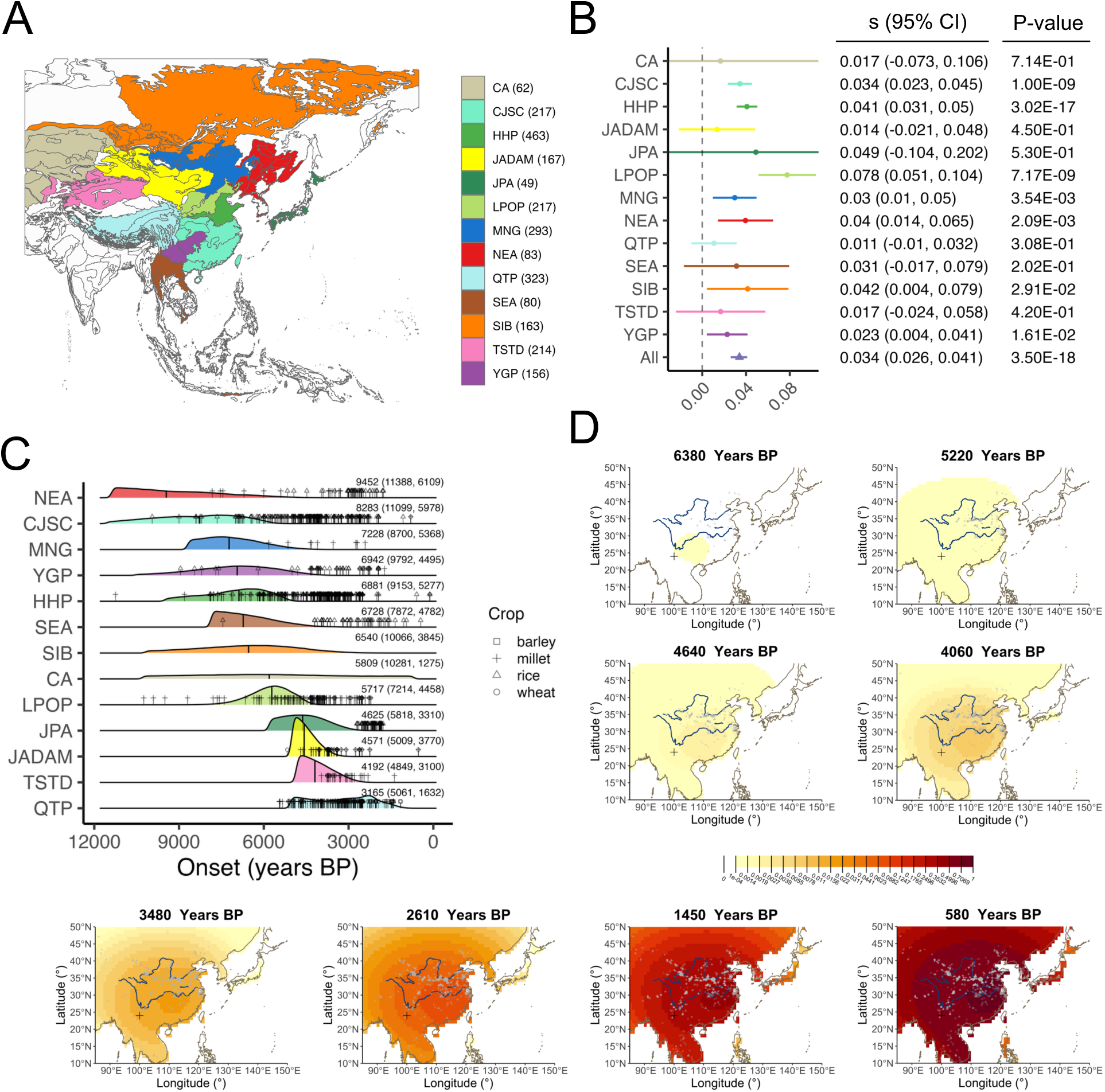
**Spatiotemporal dynamics of adaptive 48His allele of *ADH1B***. (A) Subdivision of East Eurasia for the regionally-stratified analysis of ancient genomes. (B) Forest plot of estimated selection coefficients for different regions of East Eurasia. Note that some of the 95% CI are cut off by the axis limits. (C) Posterior distribution of inferred changepoints from hinge regressions applied to the regionally stratified data. Medians are indicated by the vertical line. The appearance of various crop types in each geographic region are also plotted. (D) One spatiotemporal reconstruction of the diffusion of the adaptive allele with farm sites continually overlayed with first appearance. The "+" indicates the inferred origin of the adaptive allele based on the model. Region abbreviations: Central Asia (CA), Chang Jiang & Southern Coastal China & Sichuan Basin (CJSC), Huang He Plain (HHP), Junggar & Alashan Semi-deserts & Altai Mt. (JADAM), Japanese Archipelago (JPA), Loess Plateau & Ordos Plateau (LPOP), Mongolian Grasslands (MNG), Northeast Asia (NEA), (I) Qinghai-Tibetan Plateau (QTP), Southeast Asia (SEA), Siberia (SIB), Tianshan & Taklimakan Desert (TSTD), and Yunnan-Guizhou Plateau (YGP).

To further clarify the spatiotemporal dynamics of rs1229984, we applied an allele diffusion model that reconstructs the spread of an adaptive allele over time and space based on ancient genomes^27^, which obviates the need for geographical assignment based on hard boundaries. We implemented several versions of the model with different start times for the appearance of the adaptive allele and breakpoints for possible changes in the parameters across time. A consistent pattern across reconstructions was that the center of mass of the adaptive allele frequency was rooted in Southern China (Supplementary Videos 1-6). However, depending on the initial parameter specifications, we saw finer-scale differences for the 48His allele’s spread across East Asia. In one reconstruction, we observed that the allele frequency center mass initially fell within the middle to upper reaches of the Yangtze before advancing eastwards toward the lower Yangtze (Fig. 3D). Given the dual origins of rice farming along the Yangtze River^28^, our spatiotemporal modelling raises the possibility that the spread of this adaptive allele was more complex than a stationary diffusion from the lower Yangtze River, as was previously implicated by a study based on present-day allele frequencies in East Asia^26^. Further aDNA data from Southern China will be necessary to more precisely pinpoint the location from which the 48His allele initially expanded. What is clear is that our reconstructions do not support an origin north of the Yangtze, such as around the Jiahu site in the Central Plains, the earliest known northern agricultural site with evidence of rice farming as well as alcohol fermentation^29^.

### Association of agriculture with adaptive variants in East Asia

To formally test the relationship between agriculture and adaptive genetic variants, we constructed an environmental variable, the archaeobotanical density score (ADS), to proxy ancestral local intensities of farming (see Methods). We examined whether ancient individuals in close spatiotemporal proximity to archaeobotanical sites were more likely to carry a putatively adaptive allele compared to individuals further away (Fig. 4A). For the main selection candidates from our genome-wide scan, we tested with logistic regression for whether the ADS term significantly explains variation in genotype dosages even when accounting for shifts in ancestry and the general selection coefficient.

**Figure 4.**
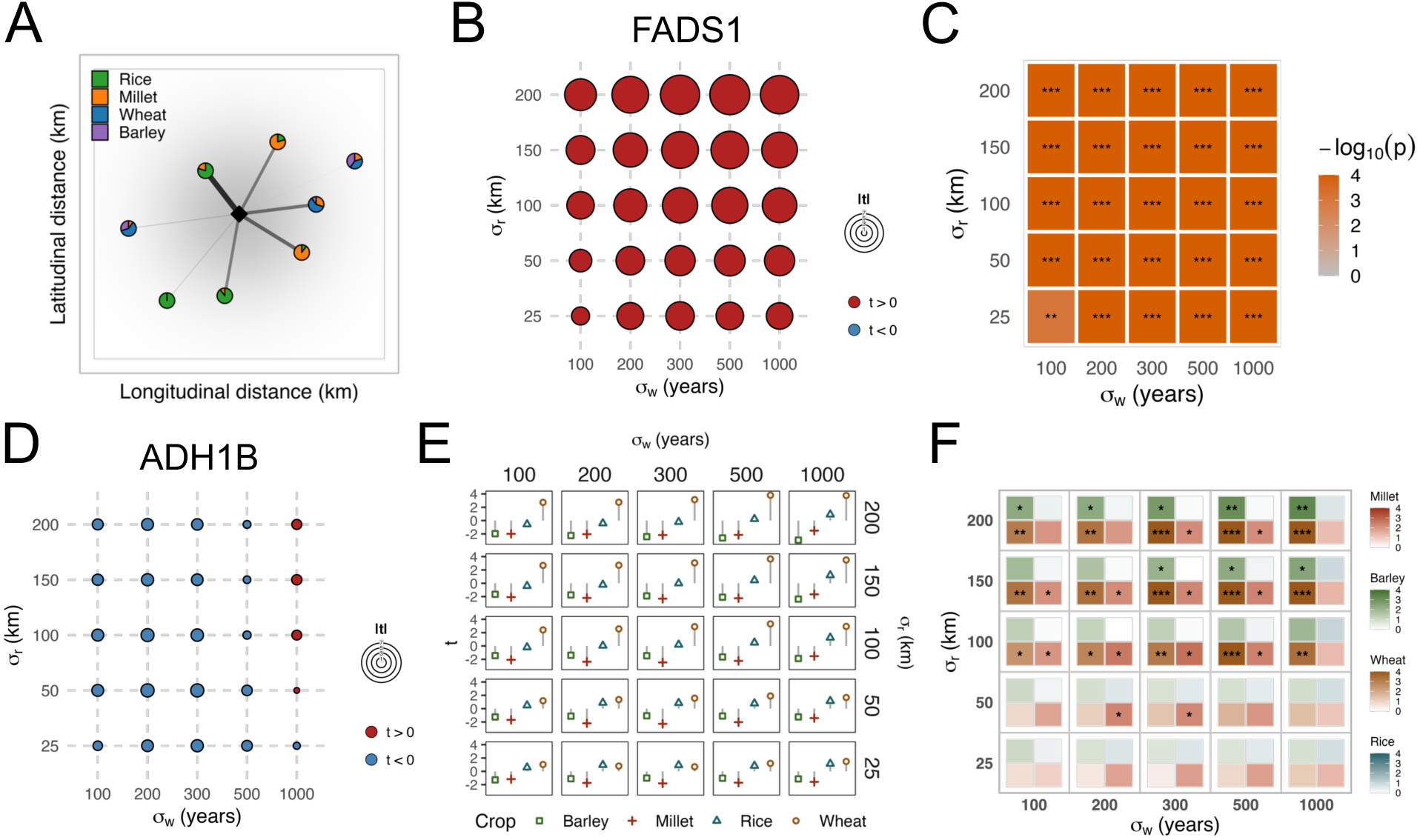
**Association of agriculture with *FADS1* and *ADH1B***. (A) Illustration of the spatial weighting of ADS for one ancient individual represented by the diamond. The pie charts represent archaeobotanical sites and their crop-specific biomass proportions. (B) Bubble plot of the association statistics for ADS from the regression of the derived allele of rs174546 across various spatial and temporal bandwidths with (C) corresponding permutation-based P-values plotted as a heatmap where P ≤ 0.01 is marked by *, P ≤ 0.001 by **, and P ≤ 1x10^-4^ by ***. (D) Bubble plot of the regression of the derived allele of rs1229984 on ADS. (E) Crop-specific ADS association statistics with the derived allele of rs1229984 and (F) a heat map of their permutation-based P-values within each block the top left cell corresponds to ADS_barley_, top right to ADS_rice_, bottom right to ADS_millet_, and bottom left to ADS_wheat_.

We observed the strongest association between ADS and the derived allele of rs174546 in *FADS1* (Fig 4B), in which under nearly all the bandwidth combinations tested the test statistic exceeded any under the null distribution based on 10,000 permutations (P < 1x10^-4^) (Fig. 4C). This result provides one of the most direct lines of evidence for a relationship between agriculture and selection at *FADS1*. When testing for associations with the crop-specific ADS, we observed opposing effects of millet and rice farming; millet farming was positively associated with the derived allele, whereas rice farming was negatively associated with it (Supplementary Fig. 24). One explanation for this pattern is that wetland environments favorable for rice paddy agriculture were rich in aquatic fauna, which are major dietary sources of LC-PUFAs. Aquaculture was even practiced in some communities from the early Neolithic, as rice paddies could host freshwater fish^30,31^. The positive association of the derived *FADS* haplotype with millet might be attributed to the relatively high content of short chain (SC)-PUFAs compared to other cereal grains^32^.

The 48His allele of rs1229984 in *ADH1B* did not show any connection to general farming intensity (Fig. 4D), but it was significantly associated with ADS_wheat_ at higher spatial bandwidths (Fig. 4E,F). While this result was unexpected given that the selected allele seems to have initially arose in Yangtze rice farmers, our model indicates that ADS_wheat_ explained increases in the 48His allele not captured by the *β_date_* term. Wheat agriculture may have induced stronger selection because its late introduction in a predominantly millet or rice-based context allowed for multicropping, as wheat is a winter crop whereas millet and rice are planted in the warmer months. Farmers would then have had a greater availability of cereal grains, of which the excess could be used for alcohol production. The *ALDH2* variant rs671, however, was associated with neither the general nor crop-specific ADS (Supplementary Fig. 25A,B and Supplementary Fig. 29A,B). Given the later selection onset and differences in relationship with agriculture, selection for the 504Lys allele might be related to a different set of environmental factors.

Finally, we observed associations with ADS for loci with functions not directly related to diet and metabolism. The light skin pigmentation allele of rs1800414 in *OCA2* was negatively associated with ADS (Supplementary Fig. 26A,B), which is in the opposite direction predicted by the hypothesized relationship in Europe. The transition to farming was thought to have resulted in a major reduction of dietary vitamin D, favoring less melanin in the epidermis^33^. Given the lower latitude of most of East Asia compared to Europe, the balance of selection pressures between maintaining adequate levels of folate and vitamin D might have been different across both ends of Eurasia. rs1047781 of *FUT2* also exhibited a surprising relationship with farming. The non-secretor allele appeared to have been selected against in environments with more rice agriculture (Supplementary Fig. 27C,D). Previous studies have suggested health tradeoffs of the non-secretor phenotype^34,35^, one of which is infant immunity to respiratory illness based on maternal *FUT2*genotypes mediated through breast milk^36^. Pathogens prevalent in wetland environments might have diminished the fitness advantages of the non-secretor allele, which could accord with the relatively high rates of childhood stress as indicated by linear enamel hypoplasias observed in Neolithic rice farming populations^37^. Similar to *FUT2*, the rs784133 variant of *RSPH4A* that was increasing in frequency over time had a negative association with ADS_barley_ (Supplementary Fig. 28C,D). As for the other selection candidates, they lacked consistent and significant associations with any ADS (Supplementary Fig. 25 and Fig. 29).

## Discussion

By analyzing some of the largest datasets of ancient human genomes and archaeobotanical sites from East Asia, we have provided deeper insights into the evolutionary dynamics of positively selected loci in East Asia. We estimated the timing and strength of selection for eight loci with genome-wide evidence for selection, reconstructed their spatiotemporal allele frequency distributions over the East Eurasian landscape, and tested their associations with agriculture based on an environmental index variable we devised, the archaeobotanical density score (ADS). Through these analyses, we addressed various hypotheses about the adaptive response to neolithization. First, our results challenge the notion that recent adaptation to pathogen exposure was primarily driven by agricultural adoption, at least with respect to the beginnings of selective sweeps. While some loci, including *HLA* and *OAS1*, started increasing in frequency during the Neolithic, most of the selection candidates related to immunity from our genome-wide scan appeared to be rising in frequency prior to the Neolithic. Furthermore, association testing with ADS generally did not indicate intensified selection on immune loci in response to farming. In fact, the East Asian-specific *FUT2* variant that began to be selected for prior to the Holocene appeared to be selected against by an environmental factor associated with rice farming despite generally continuing to rise in frequency during the Holocene. We observed a similar pattern with the missense variant of *RSPH4A* being negatively associated with barley agriculture. In contrast, the derived *FADS* haplotype, which first began rising in frequency in Upper Paleolithic hunter gatherers, exhibited a strong, positive association with ADS, indicating that the selective pressure of increased plant consumption grew with the adoption of farming. We thus demonstrate that the timing of a selective sweep by itself is neither able to implicate nor rule out agriculture’s role as a selection pressure. Selection acted on some loci from the Upper Paleolithic and continued into the Neolithic and in some cases was either amplified or lessened with neolithization.

Second, our work both supports and complicates the rice farming hypothesis that addresses the evolution of rs1229984 at *ADH1B*^26^. Through spatiotemporal allele frequency diffusion modelling, we found evidence for a southern origin along the Yangtze River, indicating an initial connection with rice agriculture. However, our ADS regression analysis showed that wheat rather than rice farming significantly explains some of the unexplained variance for the adaptive allele in our data. Based on these results, we can at least conclude that the selection pressure for the 48His missense variant of *ADH1B* was not exclusively related to rice itself; likely what mattered more was the local availability of grains for alcohol production. Third, our study adds nuance to narratives about convergent evolution across western and eastern Eurasia. The past decade of genetic studies of human adaptation has shown the targets of selection broadly being shared across diverse populations in Eurasia, which has been interpreted as representing a common response to a similar set of selective pressures brought on by neolithization^5,6,38^. Indeed, we found only a limited number of East Asian-specific loci that were under selection within the Neolithic period, such as *ALDH2*.

However, even though selection generally favored the same allele—or at least variant with respect to phenotypic direction—in East Asians and West Eurasians, aspects of the selection regime differed for several loci. For example, selection favoring the derived *FADS* haplotype and the non-secretor allele of *FUT2* appears to have begun earlier in East Asians compared to West Eurasians. Even within East Eurasia, we observed heterogeneity in the strength and timing of selection for the missense variant *ADH1B*\**48His,* underscoring the importance of environmental heterogeneity in shaping the landscape of natural selection.

Most of our selection candidates had functions clearly related to classic pathways under adaptation, namely immunity, metabolism, and skin pigmentation, but the fitness mechanism of the strongest signal in our selection scan at *ADH1B* remains unknown. The aldehyde hypothesis, which offers a possible explanation for the East Asian selection signal at *ALDH2,* proposes that elevated aldehyde levels protected against pathogens such as *Mycobacterium tuberculosis*^39–41^. However, heterozygous and homozygous carriers of the adaptive 48His allele who do not carry the *ALDH2* 504Lys variant do not exhibit high levels of acetaldehyde in the blood following ingestion of ethanol^42^, leaving selection at *ADH1B* still without adequate explanation. We propose a new hypothesis uniting several lines of evidence from epidemiology and population genetics. Vitamin B_12_ plays critical roles in DNA synthesis and red blood cell production, and pregnancies deficient in this nutrient are at heightened risk for several complications^43^. Randomized control trials have shown that moderate amounts of alcohol significantly reduce vitamin B_12_ levels in subjects^44–46^. Thus, *ADH1B*\**48His* could have provided a fitness advantage by mitigating alcohol’s inhibitory effects on vitamin B_12_ absorption. As vitamin B_12_ can only be obtained from animal-derived foods, this nutrient became scarcer as populations increasingly relied on crops for subsistence.

Concomitantly, pastoralism might explain some of the variation in allele frequency of rs1229984 across populations globally and within East Eurasia, as dairy could have served as a buffer against vitamin B_12_ deficiency. A study of East Africans found that selection had driven up the 48His allele frequency in present-day Afroasiatic-speaking agriculturalists compared to neighboring hunter-gatherers and pastoralists^47^. Though selection at *ADH1B* has also been detected in West Eurasians^2,48^, widespread pastoralism in Europe possibly blunted the extent of the selective sweep, leaving the present-day frequency at only ∼3%. As for East Eurasia, our analyses show either weak or no selection in the Qinghai-Tibetan Plateau and regions corresponding to the very northwestern parts of China and Central Asia, areas where dairy has contributed meaningfully to the local diet since at least the Bronze Age^49–51^. The Mongolian Grasslands region, however, exhibited some evidence for selection (P = 3.54x10^-3^) despite the presence of pastoralism there since 5000 BP^52^. While this observation disagrees with our hypothesis, other factors influencing the availability of this nutrient might explain this discrepancy. Vitamin B_12_ concentrations in milk vary across species; goat milk, which was consumed early on the East Eurasian Steppe^52^, for example, has very low levels^53^. In addition, the processing of milk into yogurt or cheese generally results in a loss of absolute vitamin B_12_ content, though this can depend on the specific production techniques and bacterial cultures present^54^. According to our hypothesis, the distribution of 48His allele frequency across populations should reflect the balance between historical local availability of animal-derived products and alcohol, as well as the bioavailable vitamin B_12_ and ethanol content of the kinds of food and drink available at a given locale. These quantities over time and space are generally not well characterized, necessitating further collection of archaeological dietary data to test the role of vitamin B_12_ as a selection pressure.

To conclude, our study of positive selection with ancient genomes enriches our knowledge of the evolutionary histories of major adaptive loci in East Asians and advances our understanding of the adaptive response to neolithization. Our interdisciplinary approach of jointly analyzing ancient genomic data with archaeobotanical data represents progress toward linking genomic signals of selections with environmental pressures. The gene-farming associations we find also underscore the role of balancing selection via fluctuating selection. In the vein of our study and a previous one investigating the association between dairying and the lactase persistence allele in Europe^55^, the field’s understanding of human adaptation will benefit greatly from future work that models spatiotemporal data from both ancient genomes and the environment.

## Methods

### Preparation of database

We gathered genome-wide data from ancient humans from 75 published studies^56–130^ and unpublished genomes sampled from China, for which demographic characterization will be described in forthcoming studies, and merged them with present-day East Eurasian individuals from the HGDP^131^ belonging to the following populations: Cambodian, Dai, Daur, Han, Hezhen, Japanese, Lahu, Miao, Mongolian, Naxi, Northern Han, Oroqen, She, Tu, Tujia, Uyghur, Xibo, Yakut, and Yi. If available, we started from the BAM files uploaded by the authors of publications that released data generated from outside our lab. These files were checked for whether DNA end damage masking or trimming was already performed by using PMDtools. For the majority of papers, the BAM file reads were uploaded without such treatment, so we masked bases at the ends of reads that could potentially experience deamination events based on the protocol used for ancient sequences that we have generated. For non-UDG single strand (SS) or double strand (DS) libraries we masked 5 bp from both ends, for UDG SS libraries 1 bp from the 5’ end and 2 bp from the 3’ end, and for UDG DS libraries 2 bp from both ends. Regarding contamination, we referred to the original publication’s checks for contamination and did not include individuals that were deemed unreliable by that publication. However, if publications only provided FASTQ files, we mapped to reference genome hg19 following the same protocol used for ancient genomes sequenced from our lab^115^. For individuals present in the AADR database (v62.0)^132^, we used the provided sample annotation information (e.g., date, geographic coordinates, and library style), which we also checked based on information provided in the original publications, and for all other individuals we extracted the relevant annotations directly from the papers. These metadata were synthesized into one database (Supplementary Table 1).

### Pseudohaploid genotyping of ancient genomes

We obtained pseudohaploid genotypes at 1240k sites using pileupCaller with the *randomHaploid* flag after running samtools mpileup with the arguments *-R -B -q30 -Q30* on the masked BAM files.

### Kinship-based filtering to remove close relatives and duplicates

Using READv2 (v2.01)^133^, we identified close familial related or duplicate individuals in our dataset up to the third degree of relatedness. We used pseudohaploid genotype data for the ancient samples at sites restricted to 1240k array autosomal SNPs, which were filtered for common variants (MAF>0.05). Given resource constraints of our compute environment, we ran READv2 in batches of 200 individuals with a step size of 100 wherein individuals were ordered by radiocarbon date. After running READv2, we obtained the inferred pairwise relationships based on expected kinship coefficients and applied a pruning procedure to avoid unnecessarily removing more samples than necessary while still removing closely related individuals. For individuals identified as duplicates or monozygotic twins, we took a greedy approach by pruning out the low coverage sample in all pairs. For first through third degree pairs, we prioritized pruning out the lower coverage sample for individuals with the highest number of putative connections. Based on this pruning strategy we removed 349 individuals from the database.

### Imputation of ancient genomes

We performed phasing and imputation with GLIMPSE2 (v2.0.0)^134^ on our dataset of ancient individuals using a reference panel of 2504 high coverage WGS individuals from the 1000 Genomes Project^135^, which we lifted over from reference build hg38 to hg19 using the BCFtools/liftover tool^136^. We filtered the 1000 Genomes data for biallelic SNPs and then converted used GLIMPSE2_split_reference to make it into the reference panel format. We generated genotype likelihoods for 1000 Genomes sites from the masked BAM files of the individuals to be imputed with bcftools (v1.19) mpileup (*-B -I -a ’FORMAT/DP’*) based on default mapping and base quality thresholds, followed by bcftools call ( *-Aim -C alleles*). After performing imputation with GLIMPSE2_phase in parallel on different parts of the genome, we put back together these pieces with GLIMPSE2_ligate. We extracted genotypes for 1240k capture sites from the imputed VCF with bcftools view and generated a version for the main selection scan that was based on biallelic SNPs in 1000 Genomes with MAF>0.01. For evaluating the individual level genotype accuracy of samples, we used the concordance tool provided in GLIMPSE2 with default parameters.

### Principal components analysis

Using smartPCA (version 16000)^137^, we projected the imputed ancient samples onto PC space generated from HGDP participants that were not from Africa based on LD-pruned 1240K sites. For the primary selection scan, we used the PCA coordinates based on imputed genotype data as the projected positions onto PC space should not only reflect genuine genetic relationships among individuals but also potential biases induced by imputation or technical sources of variation due to methodological differences across studies.

### Simulations to quantify imputation accuracy

To help determine the appropriate coverage threshold for inclusion of samples for our time series analysis, we performed two sets of simulation experiments based on real data. First, we examined the accuracy of using imputed low coverage data for calculating allele frequencies in the Yakut population included in HGDP. We downloaded aligned read data in CRAM format for 23 Yakut individuals and randomly downsampled (using samtools as described above) these CRAM files according to the observed distribution of coverages at 1240k sites of our original dataset of ancient genomes. For each individual, we downsampled five times based on randomly selected coverages (without replacement), generating a total of 115 downsampled diploid genomes. With these downsampled individuals, we imputed using an unrelated panel of 1000 Genomes high coverage reference panel mapped to the hg38 human reference, since the HGDP reads were also based on hg38. Finally, we calculated the allele frequencies at 1240k sites under various coverage exclusion thresholds ranging from 0.01x to 1.2x and calculated the difference between the true allele frequency in Yakut (as calculated from the original high coverage data) and imputed downsampled frequency. To investigate reference panel bias from imputation, we also compared the allele frequency based on imputed downsampled Yakut genomes to the frequency in all 1000 Genomes East Asian individuals which does not include any Siberian population.

For our second experiment, we simulated the accuracy of estimating the selection coefficient based on logistic regression using the imputed >2x coverage capture samples as a validation set. We considered analysis on the imputed genotypes at this threshold sufficient based on the individual sample genotype accuracy downsampling experiments we conducted as well as previous reports that have suggested 2x coverage as adequate for demographic analyses^138^. We downsampled these higher coverage individuals according to the distribution of our original dataset and imputed them. We calculated selection coefficients based on the main logistic regression model described below for the 1240k sites based on different exclusion thresholds ranging from 0.1x to 1x, which were then used to quantify the deviation from the selection coefficients of corresponding SNPs calculated from the original >2x dataset.

### Genome-wide logistic regression selection scan

The basis of our selection scan is to identify SNPs that changed in frequency over time more rapidly than expected, compared to the rest of the genome. We performed logistic regression for each individual SNP to predict allele frequency as a function of the radiocarbon date of the sample while controlling for covariates such as genomic PCs and coverage, to avoid false signals of selection due to confounding. The input for the dependent variable were the imputed genotype dosages from GLIMPSE2 scaled down by one half. For each SNP *i* across the genome, the following fixed effects model was applied:

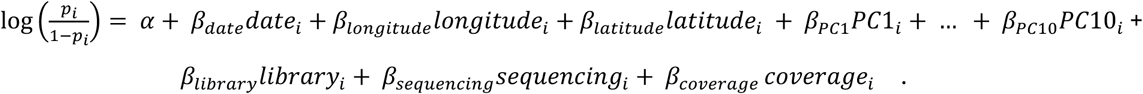

The library style variable could be one of {DS.plus, DS.minus, SS.plus, SS.minus, SS/DS.minus, SS/DS.plus, HGDP}, and we subsumed the half or partial UDG treatment samples into the full UDG treatment group. For the sequencing variable, samples could be categorized as {shotgun, capture, exome}, with capture encompassing both 1240k and Twist platforms. To obtain a P-value for the date term, which represents the signal of selection, we performed a likelihood ratio test in R with the base anova function comparing the full model with and without *β_date_*. We ran this model on all imputed SNPs for both 40,000 and 12,000 BP transects and provide the summary statistics (Supplementary Data 1 and 2).

Since *β_date_* reflects the rate of change in allele frequency over time (in units of years) independent of other variables in our model, we were able to estimate the selection coefficient for SNP *i* by multiplying *β_date_* by a generation time of 28 years^139^ and then doubling this value to reflect the difference in fitness between homozygotes with and without the selected allele in one generation.

### Estimation of allele frequency rise onset

We estimated the start time of an allele’s rise (t_on_) and obtained 95% credible intervals (CrI) by performing a Bayesian changepoint analysis with the R package mcp. With this approach, we specifically identified the time at which *β_date_* changes from zero to a non-zero slope, modelling the corresponding frequency trajectory as a hinge. For variants that show evidence of selection, we interpret t_on_ as the start of selection. We specified a normal prior, N(-5,1), truncated at -3 for the slope intercept, as we imposed upon the model an assumption of selection acting on a given variant from some low frequency. In terms of allele frequency, this prior translates to a mean onset frequency of 0.67%, with ± 2 SD corresponding to 0.09% and the ceiling of 4.7%. Given the limitations of the mcp package, the model was based on a logistic regression of hard-call genotypes, rather than the genotype dosages as used in the selection scan, on sample date without control for other covariates. For MCMC, we ran 4 chains with 20,000 iterations and discarded the first 1500 iterations for burn in, and we designated the median of the posterior distribution as t_on_ with the corresponding CrI based on the smallest bounds capturing 95% of the distribution.

### Regionally-stratified time series analysis

To investigate potential differences in selection timing and strength across East Eurasia for adaptive alleles, we divided East Eurasia into different geographic zones. We used ecological regions that were proposed for the purposes of conservation biology^140^, which served as a heuristic for capturing spatial differences across East Eurasia in climate, agriculture, and population history. Given the paucity of data in some of the ecoregions, especially in southern East Asia and southeast Asia, we amalgamated ecoregions into larger regions as shown in Supplementary Table 2, resulting in 13 regions for analysis. For the stratified logistic regression models, we used robust generalized linear model with a binomial link function to downweight data points with outsized influence over the regression estimates. The temporal coverage was especially sparse at older timepoints in some regions, resulting in unstable estimates for the slopes driven by some high leverage points. We implemented the following logistic regression independently for each region in East Eurasia with the glmrob function from the robustbase R package^141^:

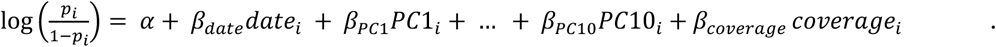

Based on the estimates of *β_date_* from the 13 regions, we conducted a random effects meta-analysis with the rme package in R with the metafor package^142^.

### Estimation of selection coefficients and allele frequency trajectories with CLUES

We used CLUES2^143^, an HMM-based approach that infers changes in allele frequency over time and the selection coefficient, to check for correspondence with our logistic regression modelling results. As CLUES2 requires an ancestral recombination graph (ARG) representative of the population being modelled as input, we utilized the published ARGs provided by Speidel et al.^144^ generated from Relate based on genome-wide joint inference of all samples from the 1000 Genomes Project. We extracted the subtrees corresponding to the East Asian populations (CHB, CHS, CDX, KHV, and JPT) from the full 1000 Genomes ARG using the SubTreesForSubpopulation module of Relate and used these subtrees for downstream analyses. To generate the .coal files, which encode the population history for subsequent selection inference with CLUES2, we re-estimated the historical effective population size of East Asians using the calibrated branch lengths from the extracted subtrees, covering all the autosomes, and assumed a per-generation mutation rate of μ = 1.25 × 10⁻⁸ and generation time of 28 years. We then sampled branch lengths from these Relate output files.

Selection inference was performed using CLUES2 based on the ancient DNA time series data and contemporary derived allele frequencies from 1000 Genomes East Asian populations. For the ancientSamps input which contains the genotype likelihoods of the ancient individuals, we transformed the genotype probabilities calculated by GLIMPSE2 into likelihoods assuming a uniform prior, following the approach of a previous study of West Eurasians that found that this assumption does not result in systematic biases^3^. We restricted this analysis to samples with at least 0.5x coverage. We set the time range for reconstruction of allele frequency trajectories at 42,000 years before present (tCutoff = 1500 generations). To examine differences in the strength of natural selection before and after the start of the Neolithic, we also ran CLUES2 using a two epoch scheme split at 10,000 BP (357 generation bins).

### Modelling of allele frequency diffusion in time and space

To reconstruct the allele frequency landscape of adaptive variants over time and space, we used a method suitable for genotype data from ancient individuals^27^. The underlying model for this method parameterizes the spread of an allele over a landscape from the inferred origin point of the beneficial mutation with migration from the center mass of the allele source with advection terms, and the selection coefficient. Two sets of these parameters were estimated before and after 5000 years BP, as intensification of genetic exchanges between the northern and southern populations in China around this timepoint have been observed in previous studies. We made a change to the published R program for doing the maximum likelihood estimation by allowing it to use genotype dosages as inputs, as that was the output of GLIMPSE2, whereas the original version only accepted either hard call genotypes or genotype likelihoods. Given that the original publication developed the software with a focus on studying West Eurasia, we had to make adjustments to the original code to adhere to the geographic boundaries of East Eurasia (e.g., setting the annealing points for inferring the beneficial allele origin). To incorporate samples from Japan and Taiwan, we imposed land bridges to mainland East Asia by manually setting those grid points that were underwater to above sea level (Supplementary Fig. 16). For the age of alleles, we relied on estimates from the public database Atlas of Variant Age^145^. For rs1229984, since the selected allele was not the alternate allele, we used an estimate from one publication studying *ADH1B*48His*^26^ and we also ran the model assuming 10,000 BP to represent the start of the Neolithic. We incorporated 1000 Genomes East Asian individuals in addition to the HGDP data to provide more data for present day. Animations for the spatiotemporal allele frequency distributions of the main selection candidates are provided in Supplementary Videos 1-26.

### Assembly of archaeobotanical database

We compiled an archaeobotanical dataset comprising 1220 observations from 862 archaeological sites across East Eurasia (China, 1020; Cambodia, 10; Indonesia, 1; Japan, 77; Laos, 2; Malaysia, 2; Myanmar, 1; Pakistan, 7; Philippines, 2; South Korea, 63; Thailand, 22; Vietnam, 13). In this database, each site was counted as one spatial unit, while multiple records from the same site representing different chronological phases were treated as distinct observations. The dataset includes macrofossil counts of cereal crops (foxtail millet, broomcorn millet, rice, wheat, and barley) spanning a time range from approximately 10,000 to 2000 BP (Supplementary Fig. 18-21). The data were collected from relevant literature and published archaeobotanical databases^61,146–150^ (Supplementary Table 3). Chronological data were directly extracted from the source literature. For sites with multiple dates, we defined the temporal span using the earliest and latest available ages. These dates consist of either direct radiocarbon dating of botanical remains or dates based on the associated archaeological period. For sites lacking original chronological information, we retrieved radiocarbon dates if available or other date estimates from the literature. Geographic coordinates were obtained directly from the source literature if available. For early excavations where coordinates were missing, locations were georeferenced based on site maps provided in the literature or localized to the specific county or village mentioned in official archaeological reports.

To ensure the independence of each data point, we implemented a multi-stage deduplication process. As data redundancy was exclusively identified within records from China, duplicates were identified and removed based on a combination of administrative province, site name, and chronological phase. Given that the dataset by Gong et al.^146^ lacked explicit primary references, we applied a spatial distance threshold of approximately 75 km using geographic coordinates to identify potential overlaps with the database of Fuller et al.^149^. Through this geospatial filtering, 136 suspected duplicate records were excluded. Furthermore, we performed a secondary audit of the database to remove sites where all archaeobotanical data were recorded as zero or lacked positive identification for any target taxa. To account for varying quantification methods employed by different scholars, which may introduce uncertainties in absolute seed counts, we standardized the data; where precise numerical counts were unavailable, presence/absence was denoted as "X" in Supplementary Table 3. Records based on plant microfossils were specifically marked as ‘X*’. For the Gong et al.^146^ database only rice sites were recorded and in some of the other published databases barley records were not provided. For downstream analyses, we treated the unknown values of the other crops as 0.

### Spatiotemporal association of agriculture with selected alleles

We devised an index, the archaeobotanical density score (ADS), to investigate variation among ancient individuals in their ancestral exposure to agriculture. The assumption behind ADS is that the density of archaeobotanical sites in proximity to an individual correlates with that individual’s ancestors’ exposure to selection pressures related to farming. This index was loosely inspired by a classic method in spatial point pattern analysis, Ripley’s K^151^. While in preliminary analyses we attempted to implement the bivariate spatiotemporal version of Ripley’s K, which detects deviations from random dispersion between events of two different classes in space and time, the sparsity and unevenness of our genetic and archaeobotanical databases made this approach difficult to use. Ripley’s K controls for variation in the background rate of events by parameterizing them as background intensities, and the dearth of observations in some patches of space and time in our data resulted in near zero values that resulted in extreme values of the statistic. Thus, instead of testing for a spatiotemporal association by modelling the appearance of alleles and archaeobotanical sites as distinct Poisson processes, we approached this problem in a regression framework. We tested for whether the variance in genotype dosage of ancient individuals in our dataset could be explained by ADS beyond the general selection term (*β_date_*) and ancestry. This approach should be most suitable for a scenario in which the selection pressure appears heterogeneously across space and time, generating variance in allele frequencies that cannot be modelled by a uniform selection coefficient. A non-significant ADS association therefore does not necessarily contradict the possibility that an environmental variable related to farming drove selection as modelled by the overall increase in allele frequency over time.

First, we calculated the ADS for each ancient individual *i* as

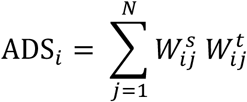

where *N* is the number of archaeobotanical sites and 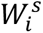 and 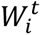 are the spatial and temporal weights of a given site relative to the *i^th^* individual. For the spatial weight, we used a standard Gaussian kernel, 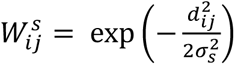, where d*_ij_* is the great circle distance between individual *i* and archaeobotanical site *j*, which was calculated using the st_distance function from the R package sf, and 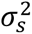 is the spatial bandwidth which modulates the rate of decay of a site’s weight with distance. We used a piecewise function based on Gaussian kernels with asymmetric rates of decay for the temporal weight because the environmental variable exerting the selection pressure necessarily precedes the expansion of the adaptive allele:

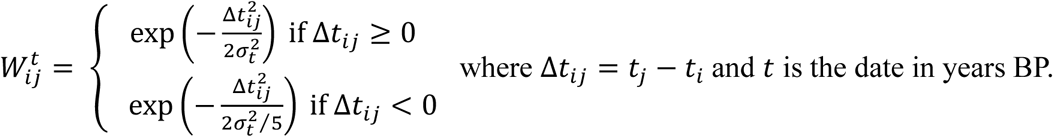

We did not entirely discount archaeobotanical sites that have dates after a given ancient sample because of error in radiocarbon dating for both the ancient samples and archaeobotanical sites. For such sites, we scaled down 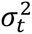 by a factor of five. Similar to how Ripley’s K is evaluated over a range of spatial and temporal windows, we calculated ADS across varying bandwidths. The Gaussian kernel adjusts the relative weighting of each site in accordance with the spatial and temporal bandwidth settings. Specifically, we tested 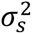 = {25, 50, 100, 150, 200} and 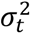 = {100, 200, 300, 500, 1000} in units of km and years, respectively.

Leveraging the information of grain counts for specific crops, we further developed ADS to represent intensities for rice, millet, barley, and wheat. The crop-specific ADS was calculated by incorporating an additional term, *W_c_*, that adjusts the weight of an archaeobotanical site by the relative contribution of a given crop to the site’s total biomass, serving as a crude proxy for the intensity of agriculture of a specific crop. Specifically, we calculated ADS for a given crop *c* as

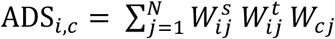

where 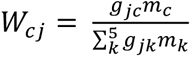 . Here, *g* refers to the grain count and *m* the thousand grain weight (TGW) of crop *c* or *k*, which we assumed to be 20 g for rice, 45 g for barley, 35 g for wheat, 2.6 g for foxtail millet, and 7.5 g for broomcorn millet. We note that the archaeobotanical literature does not appear to have a good estimate for TGW for rice, so we had to rely on modern estimates, which might overestimate the contribution of rice. Given the weight differences between foxtail and broomcorn millet, we pooled them after separately calculating the weights such that *W_millet_* = *W_foxtail_ + W_broomcorn_.* In the cases where grain counts were unavailable but the publication or database provided information about the types of cereal remains observed, we evenly split the weight of that site.

For the ADS analysis, we excluded present-day individuals given that our database has very few sites in historical times. We also excluded ancient genomes from the regions SIB and CA, as we were unable to find records of archaeobotanical sites for these regions. These filters left a sample size of 2030 individuals for the regression. We included ADS as a predictor in a logistic regression model for genotype dosage of SNP *l* that included the selection term that served as the basis for our time series selection as well as other relevant covariates such as ancestry:

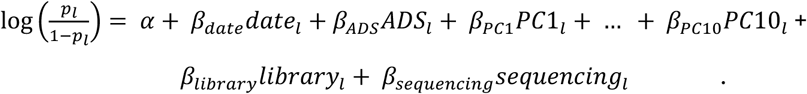

To assess the statistical significance of the ADS association, we obtained P-values for the genetic association of ADS or crop-specific ADS based on distributions of the Wald statistic from 10,000 permutations. We shuffled the residuals of ADS on the other covariates in the model in order to account for the correlation of these variables with ADS, a strategy with good performance based on one study that refers to it as the Smith procedure^152^. The ADS residuals were obtained using a general linear model (GLM) with a Tweedie distribution, which we chose given that the observed ADS distributions were right-skewed, continuous, and positive, and contained zero or practically zero values, particularly for the smaller bandwidths (Supplementary Fig. 22).

We examined the crop-specific ADS associations using the following logistic regression model that jointly incorporated the four crop scores:

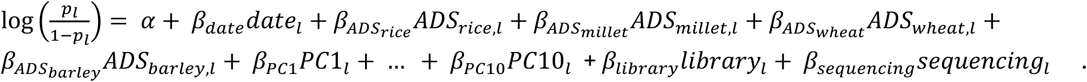

For assessing significance, we again calculated P-values based on 10,000 permutations with the Smith procedure, but rather than shuffling each crop-specific ADS independently of each other, we shuffled their residuals together as a block. To obtain these residuals, we used the same Tweedie GLM in which a crop-specific ADS was regressed over the covariates of the above model except for the scores representing the other crops.

## Data availability

The genomic data for the 236 new ancient genomes will be released in their respective publications along with accompanying archaeological information and metadata by the time of this manuscript’s final publication. Upon final publication, the data annotations for these new ancient individuals will be included in Supplementary Table 1, and the imputed genotypes for all 2236 ancient individuals used in this study will be released in a publicly accessible repository hosted on OMIX, China National Center for Bioinformation/Beijing Institute of Genomics, Chinese Academy of Sciences at https://ngdc.cncb.ac.cn/omix (accession no. PRJCA076434). Selection scan summary statistics for Supplementary Data 1 and 2 are available for download at https://doi.org/10.6084/m9.figshare.34297290 and https://doi.org/10.6084/m9.figshare.34304319, respectively. Animations for the spatiotemporal allele frequency reconstructions for the selection candidates corresponding to Supplementary Videos 1-26 are available at https://doi.org/10.6084/m9.figshare.34327725.

## Code availability

The code for running the major analyses can be found in the following GitHub repository: https://github.com/Dan-Ju/aDNA_EA_selection.

## Acknowledgments

We thank Lili Lu for providing guidance on the initial preparation of the archaeobotanical database.

## Author contributions

D.J. and Q.F. designed the study. Q.F. supervised the research project. D.J., X.F., and T.L. processed the genomic data. C.L., X.W., and M.Z. prepared the archaeobotanical database. D.J., M.Z., and T.L. curated the archaeological annotations for the ancient individuals. D.J. developed the statistical methodology. D.J. and M.Z. analyzed the data. D.J. and Q.F. discussed the analyses and the results. D.J. designed the figures. D.J. and Q.F. wrote the manuscript. C.L. and M.Z. contributed to the writing of the Methods. Q.F. and M.S. advised on the interpretation of results and critically revised the manuscript. D.J. prepared the supplementary materials. Q.F. and P.C. supervised the wet laboratory work. All authors discussed, revised, and approved the final version of the paper.

## Supplementary information

Supplementary Figures 1-32, Tables 1-3, Data 1-2, Videos 1-26, and Note can be found in the Supplementary Information file.

