## Supplement for "Ancient genomes chronicle the history of adaptive evolution in East Asia"

#### Supplementary Tables

**Supplementary Table 1:** Archaeological metadata for published East Eurasian ancient samples.
**Supplementary Table 2:** Categorization scheme of East Eurasian ecoregions into broader regions.
**Supplementary Table 3:** Archaeobotanical database of sites with evidence of cereal grains in East Eurasia.

#### Supplementary Data

**Supplementary Data 1:** Selection scan summary statistics based on a 12,000 BP time transect.
**Supplementary Data 2:** Selection scan summary statistics based on a 40,000 BP time transect.

#### Supplementary Videos

**Supplementary Video 1:** Allele frequency spatiotemporal reconstruction for rs1229984 with 1 epoch model assuming an allele age of 10,000 BP.
**Supplementary Video 2:** Allele frequency spatiotemporal reconstruction for rs1229984 with 2 epoch model assuming an allele age of 10,000 BP.
**Supplementary Video 3:** Allele frequency spatiotemporal reconstruction for rs1229984 with 1 epoch model assuming an allele age of 15,551 BP.
**Supplementary Video 4:** Allele frequency spatiotemporal reconstruction for rs1229984 with 2 epoch model assuming an allele age of 15,551 BP.
**Supplementary Video 5:** Allele frequency spatiotemporal reconstruction for rs1229984 with 1 epoch model assuming an allele age of 6048 BP.
**Supplementary Video 6:** Allele frequency spatiotemporal reconstruction for rs1229984 with 2 epoch model assuming an allele age of 6048 BP.
**Supplementary Video 7:** Allele frequency spatiotemporal reconstruction for rs671 with 1 epoch model assuming an allele age of 10,000 BP.
**Supplementary Video 8:** Allele frequency spatiotemporal reconstruction for rs671 with 2 epoch model assuming an allele age of 10,000 BP.
**Supplementary Video 9:** Allele frequency spatiotemporal reconstruction for rs671 with 1 epoch model assuming an allele age of 4034 BP.

**Supplementary Video 10:** Allele frequency spatiotemporal reconstruction for rs1800414 with 1 epoch model assuming an allele age of 24,659 BP.
**Supplementary Video 11:** Allele frequency spatiotemporal reconstruction for rs1800414 with 2 epoch model assuming an allele age of 24,659 BP.
**Supplementary Video 12:** Allele frequency spatiotemporal reconstruction for rs174546 with 1 epoch model assuming an allele age of 17,374 BP.
**Supplementary Video 13:** Allele frequency spatiotemporal reconstruction for rs174546 with 2 epoch model assuming an allele age of 17,374 BP.
**Supplementary Video 14:** Allele frequency spatiotemporal reconstruction for rs79230913 with 1 epoch model assuming an allele age of 6661 BP.
**Supplementary Video 15:** Allele frequency spatiotemporal reconstruction for rs79230913 with 2 epoch model assuming an allele age of 6661 BP.
**Supplementary Video 16:** Allele frequency spatiotemporal reconstruction for rs601338 with 1 epoch model assuming an allele age of 12,000 BP.
**Supplementary Video 17:** Allele frequency spatiotemporal reconstruction for rs601338 with 2 epoch model assuming an allele age of 12,000 BP.
**Supplementary Video 18:** Allele frequency spatiotemporal reconstruction for rs1047781 with 1 epoch model assuming an allele age of 15,812 BP.
**Supplementary Video 19:** Allele frequency spatiotemporal reconstruction for rs1047781 with 2 epoch model assuming an allele age of 15,812 BP.
**Supplementary Video 20:** Allele frequency spatiotemporal reconstruction for rs1047781 with 2 epoch model assuming an allele age of 50,985 BP.
**Supplementary Video 21:** Allele frequency spatiotemporal reconstruction for rs784133 with 1 epoch model assuming an allele age of 22,760 BP.
**Supplementary Video 22:** Allele frequency spatiotemporal reconstruction for rs784133 with 2 epoch model assuming an allele age of 22,760 BP.
**Supplementary Video 23:** Allele frequency spatiotemporal reconstruction for rs9258176 with 1 epoch model assuming an allele age of 7337 BP.
**Supplementary Video 24:** Allele frequency spatiotemporal reconstruction for rs9258176 with 2 epoch model assuming an allele age of 7337 BP.
**Supplementary Video 25:** Allele frequency spatiotemporal reconstruction for rs6766450 with 1 epoch model assuming an allele age of 9631 BP.
**Supplementary Video 26:** Allele frequency spatiotemporal reconstruction for rs6766450 with 2 epoch model assuming an allele age of 9631 BP.

#### Supplementary Information

**Supplementary Note 1:** Evaluation of imputation accuracy in ancient genomes from East Asia.

### Supplementary Figures

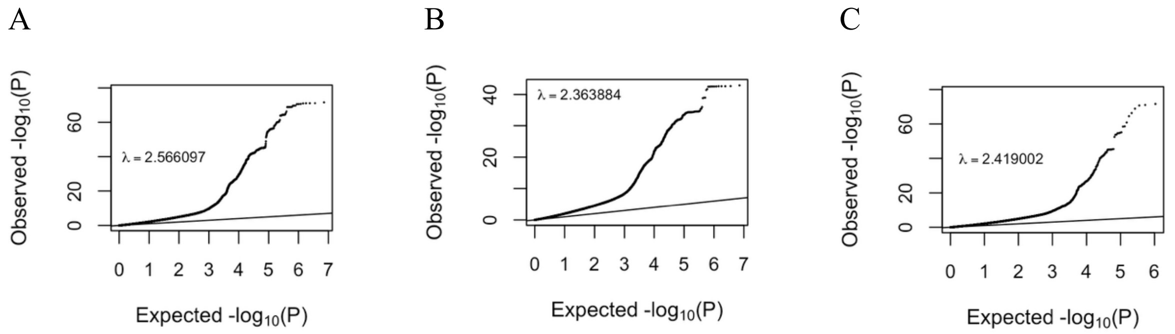


**Figure 1.** QQ plots for distribution of date term P-values for (A) imputed sites over 12,000 years, (B) imputed sites over 40,000 years, and (C) 1240k sites over 12,000 years.

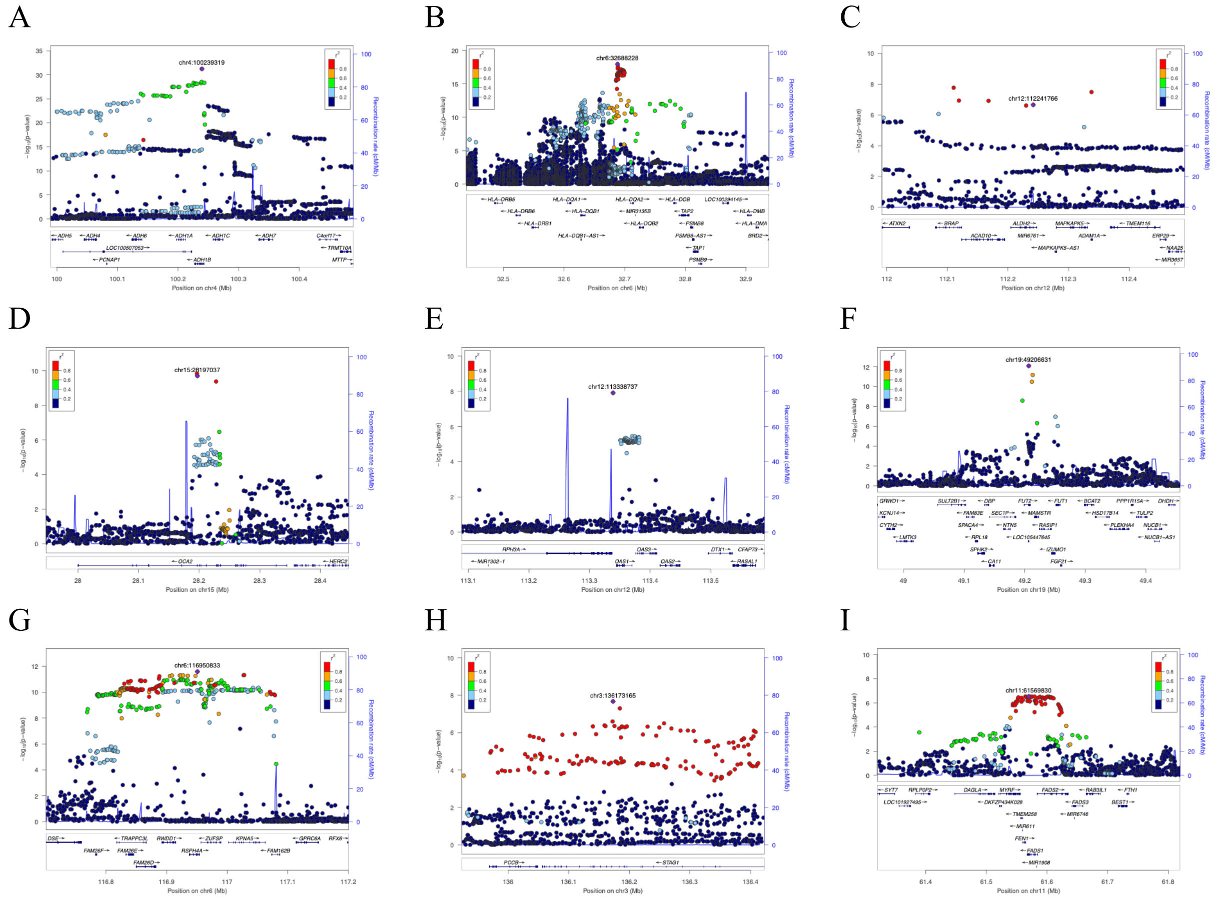


**Figure 2.** LocusZoom plots of the eight genome-wide significant loci (A-I) from the main time series selection scan. For the locus covering *ALDH2*, we used rs671 as the focal variant despite not being the most significant SNP because rs671 is a well-known missense variant with clear phenotypic effects that has also been shown in previous studies to be under selection.

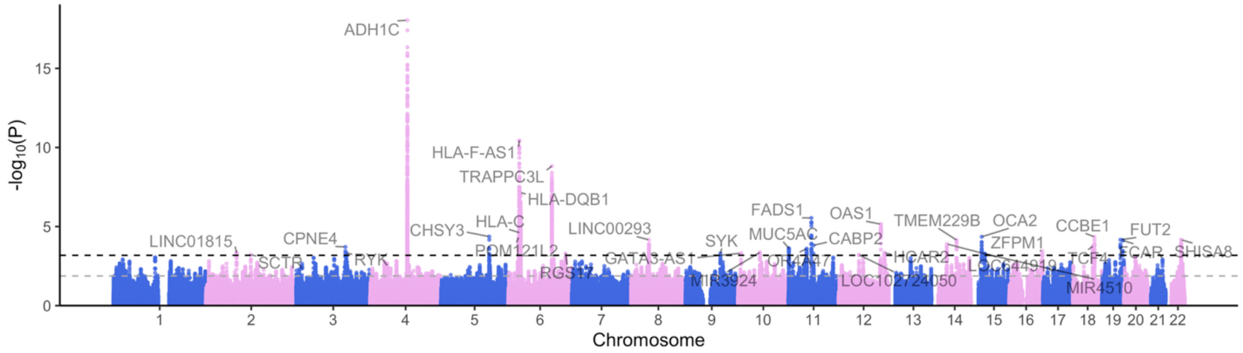


**Figure 3.** Manhattan plot of 21-SNP window averaged P-values from logistic regression selection scan analysis for all imputed sites at common SNPs in 1000 Genomes over 12,000 year period. The lines here represented empirical at the 99.95 and 99.9 percentiles.

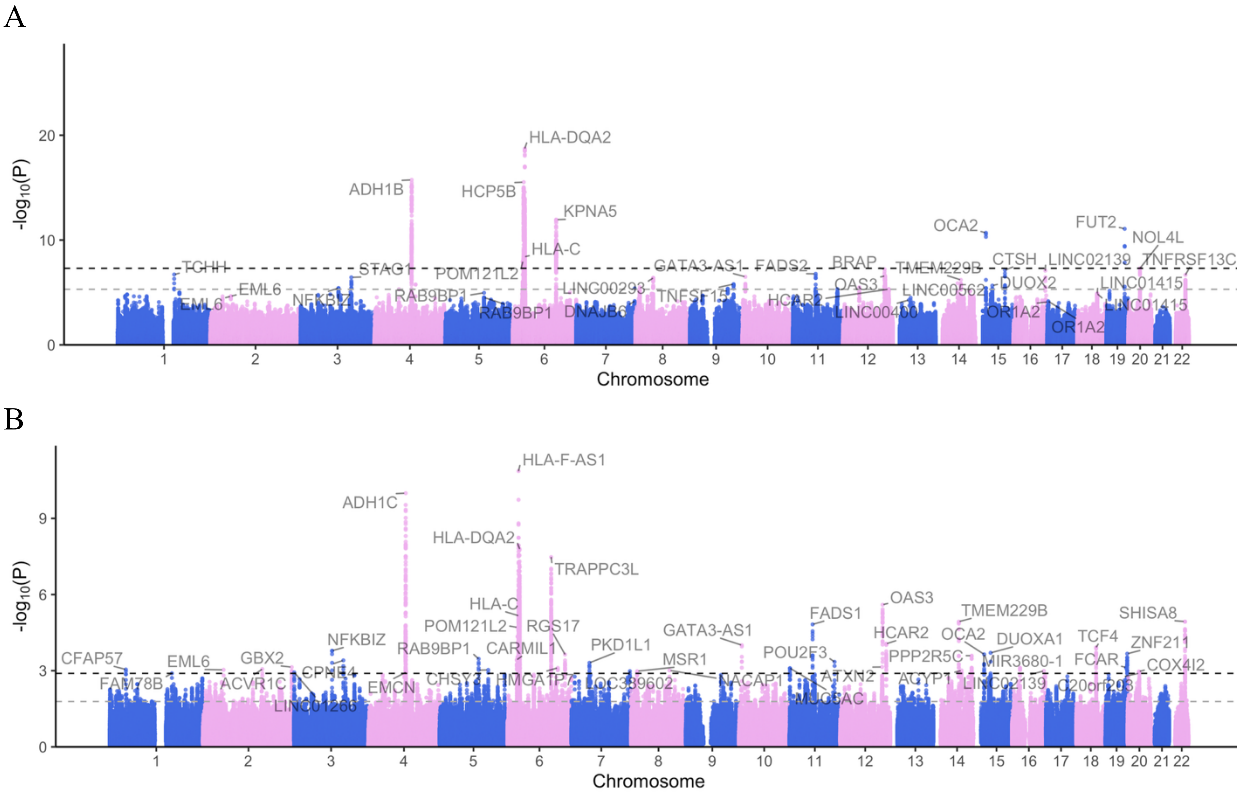


**Figure 4.** Manhattan plot of (A) individual SNP and (B) 21-SNP window averaged P-values from logistic regression selection scan analysis for all imputed common SNP sites in 1000 Genomes over 40,000 year period (N=2469).

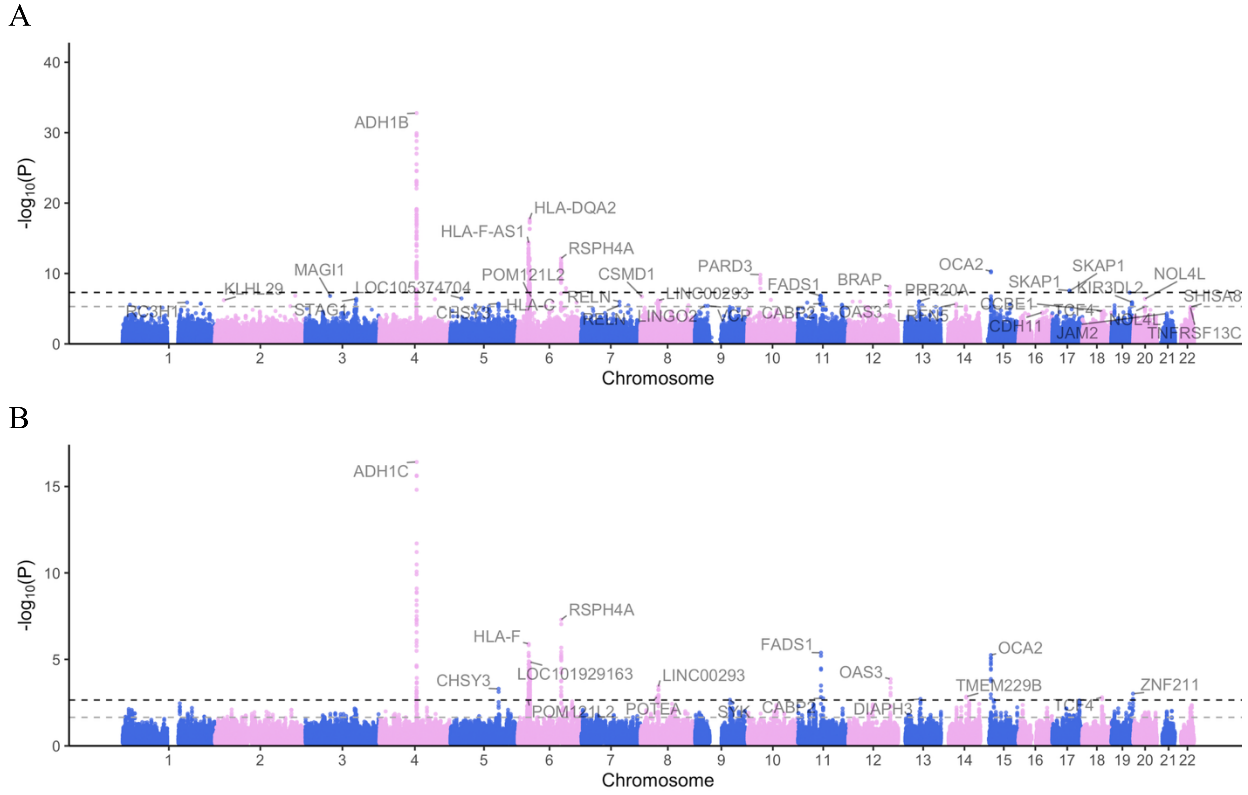


**Figure 5.** Manhattan plot of (A) individual SNP P-values and (B) averaged P-values of 21-SNP windows from logistic regression selection scan analysis based on 1240K sites only.

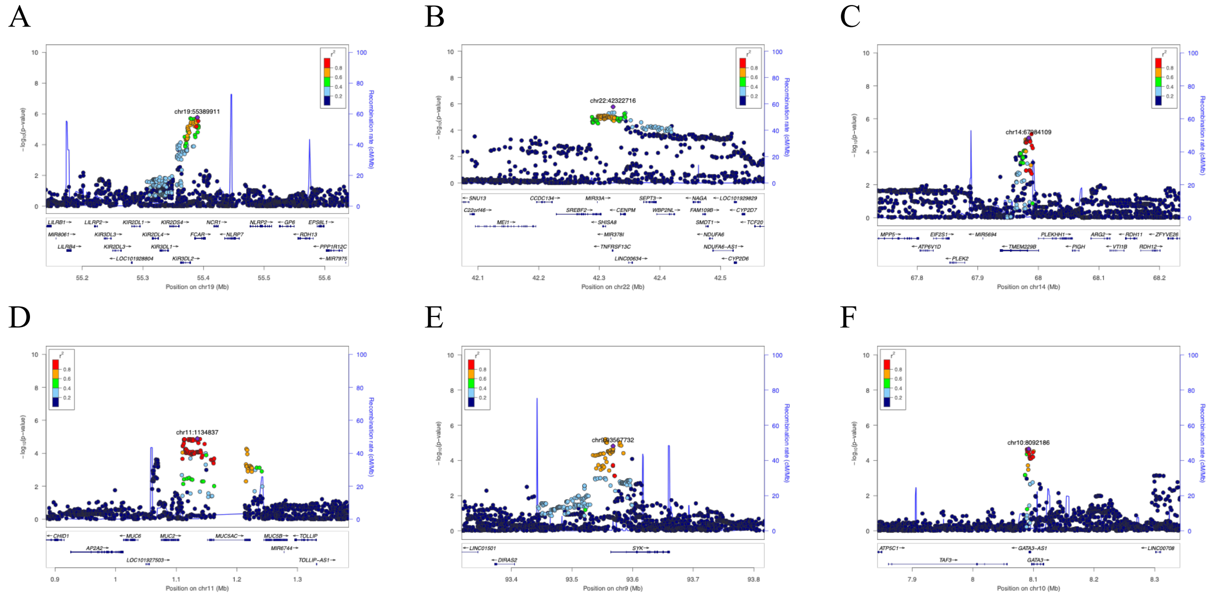

**Figure 6.** (A-F) LocusZoom plots based on outlier genomic windows overlapping genes known to be involved in immunity. The overlapping or nearest genes for the top SNPs in each locus are (A) *FCAR*, (B) *TNFRSF13C*, (C) *TMEM229B*, (D) *MUC5AC*, (E) *SYK*, and (F) *GATA3*.

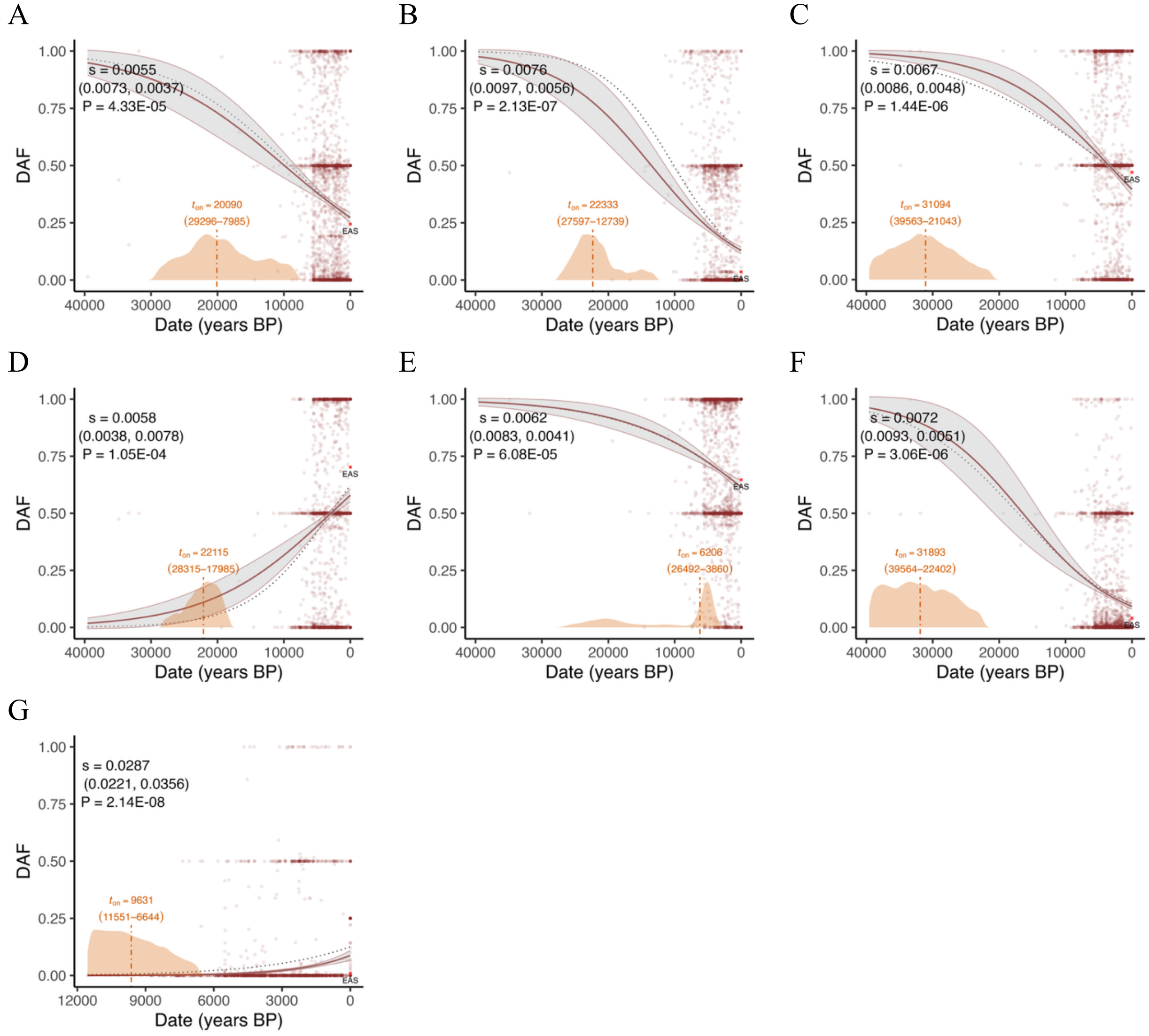


**Figure 7.** (A-F) Allele frequency trajectories for top SNPs from outlier genomic windows containing known immune genes: (A) rs56216868 (*FCAR*), (B) rs77874543 (*TNFRSF13C*), (C) rs7144624 (*TMEM229B*), (D) rs12791023 (*MUC5AC*), (E) rs1333632 (*SYK*), and (F) rs1271899 (*GATA3*), and (G) rs6766450 (*STAG1*). Note that for rs56216868 we assumed the alternate allele was the derived allele because we were not able to find an ancestral allele assignment.

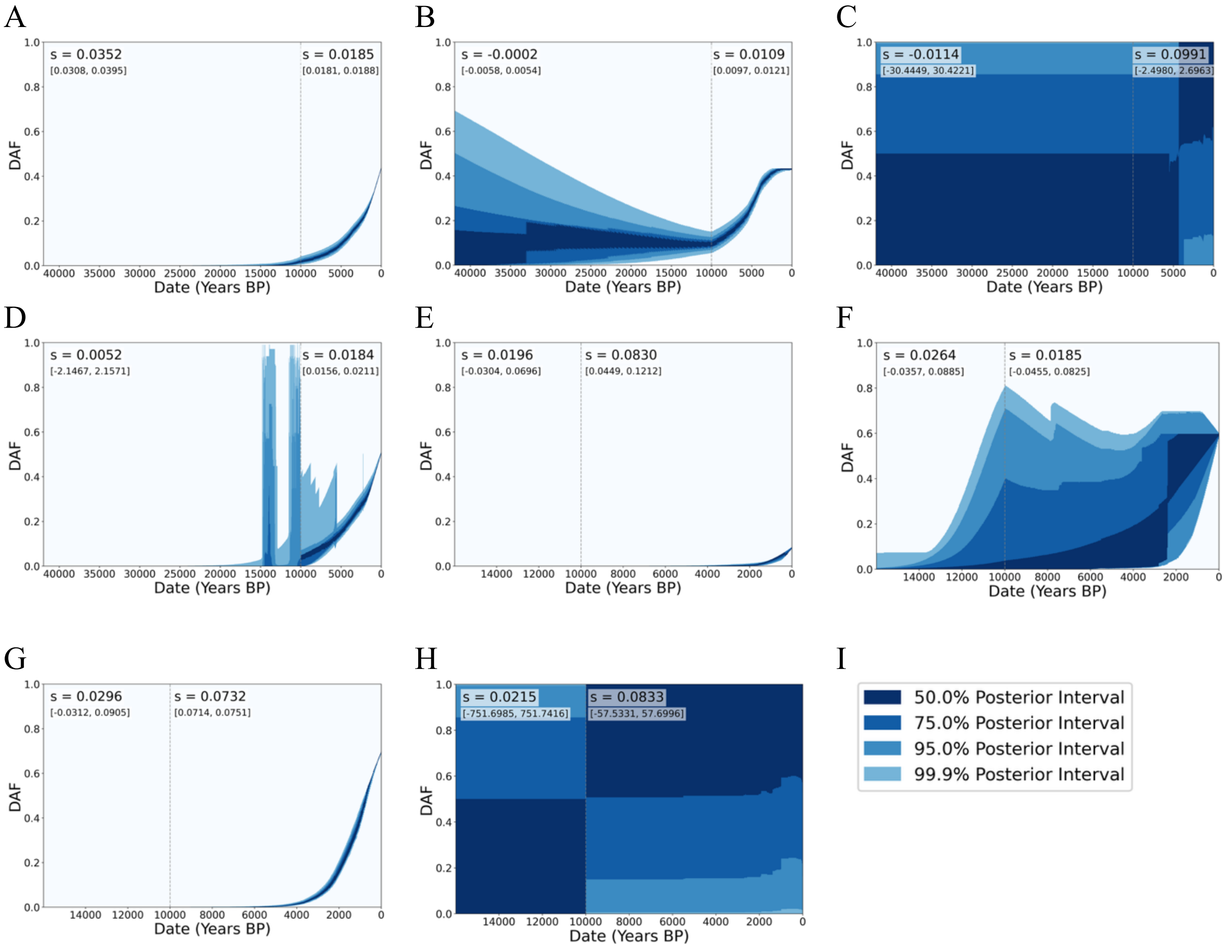


**Figure 8.** Allele frequency trajectories inferred from CLUES2 for (A) rs1047781 at *FUT2* (B) rs174546 at *FADS1*, (C) rs9258176 *HLA-DQA2*, (D) rs784133 at *RSPH4A*, (E) rs79230913 near *OAS1*, (F) rs1800414 at *OCA2*, (G) rs1229984 at *ADH1B*, and (H) rs671 at *ALDH2* using a two epoch model with the boundary demarcated at the dash line representing a rough start time for the Neolithic in East Asia. (I) Color key for different posterior intervals used in the CLUES2 frequency reconstructions.

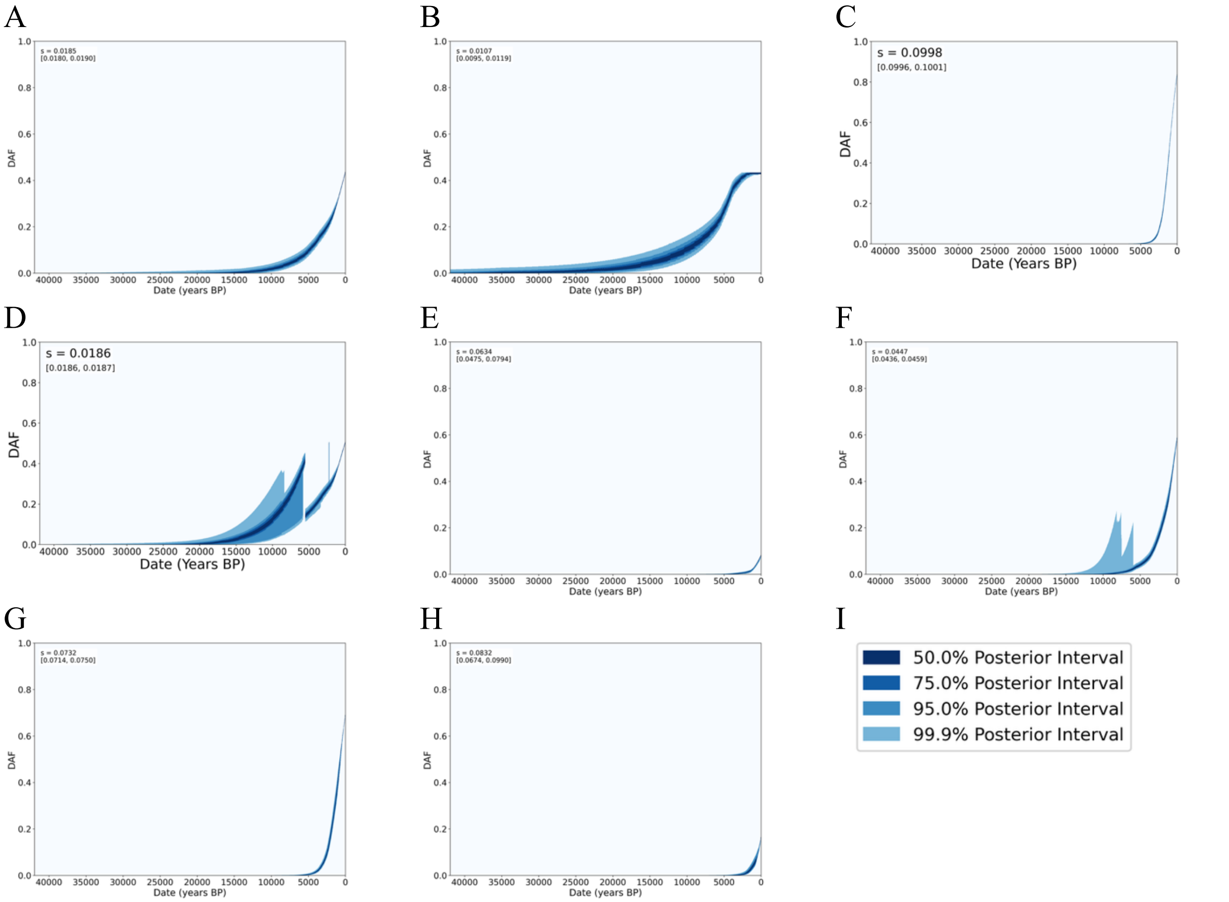


**Figure 9.** Allele frequency trajectories inferred from CLUES2 for (A) rs1047781 at *FUT2* (B) rs174546 at *FADS1*, (C) rs9258176 *HLA-DQA2*, (D) rs784133 at *RSPH4A*, (E) rs79230913 near *OAS1*, (F) rs1800414 at *OCA2*, (G) rs1229984 at *ADH1B*, and (H) rs671 at *ALDH2* using a one epoch model. (I) Color key for different posterior intervals used in the CLUES2 frequency reconstructions.

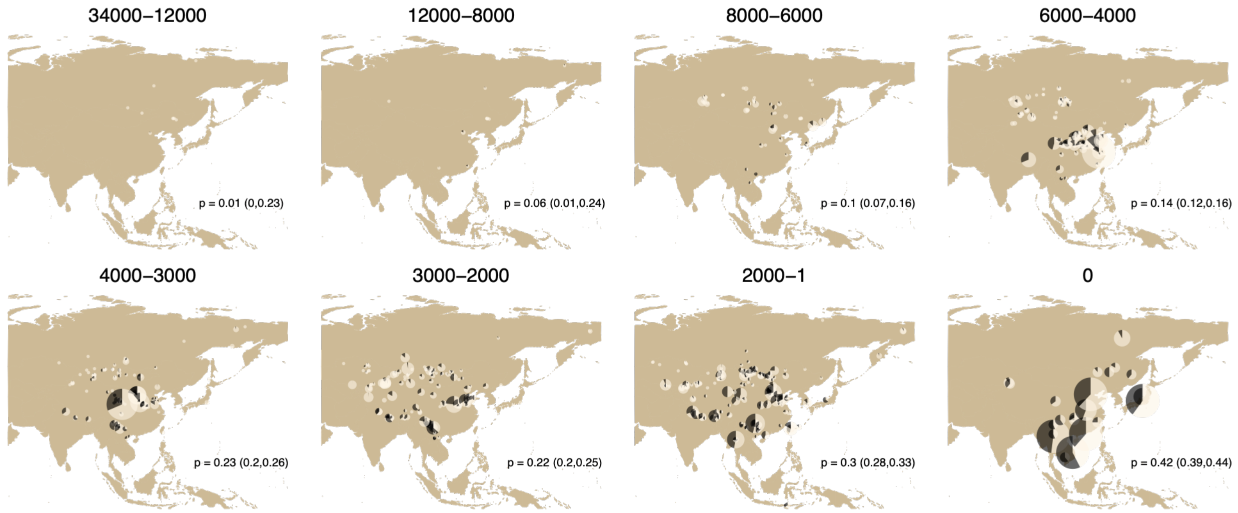


**Figure 10.** Derived allele frequency map based on imputed genotype dosages for the common East Asian missense variant rs1047781 (*FUT2*).

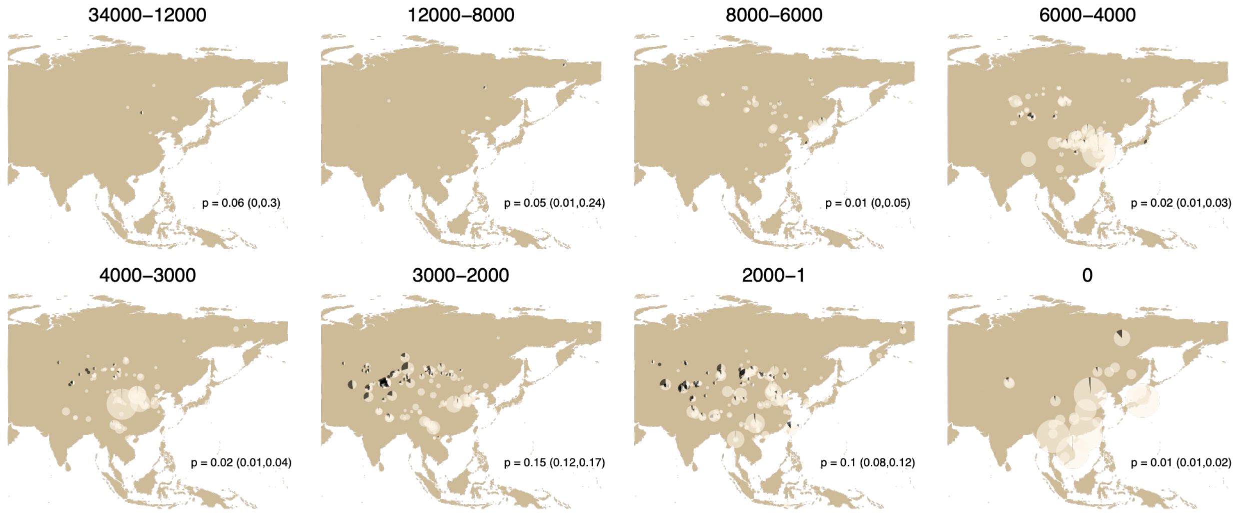


**Figure 11.** Derived allele frequency map based on imputed genotype dosages for the common West Eurasian truncation variant rs601338 (*FUT2*).

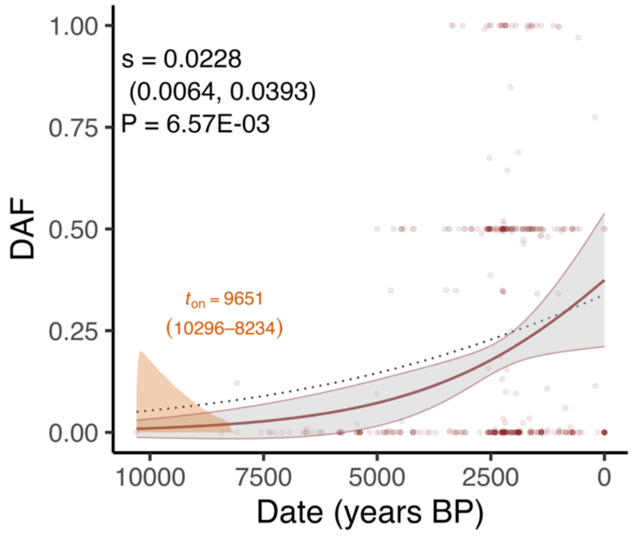


**Figure 12.** Allele frequency trajectory in the northwestern subset (corresponding to the regions of CA, JADAM, and TSTD in main Figure 3) of the genomic database.

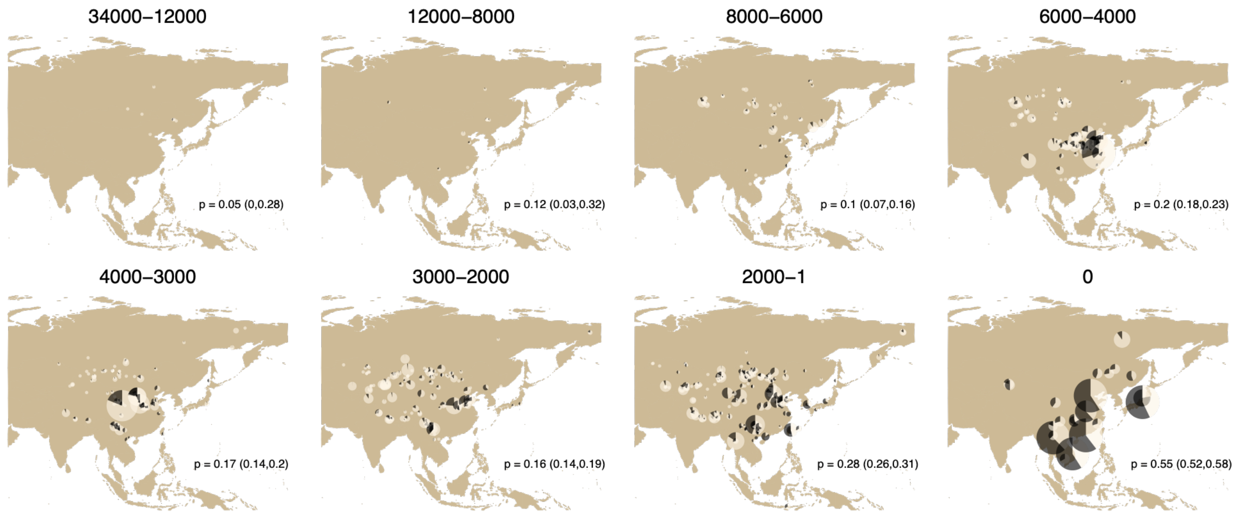


**Figure 13.** Derived allele frequency map based on imputed genotype dosages for the skin pigmentation variant rs1800414 (*OCA2*).

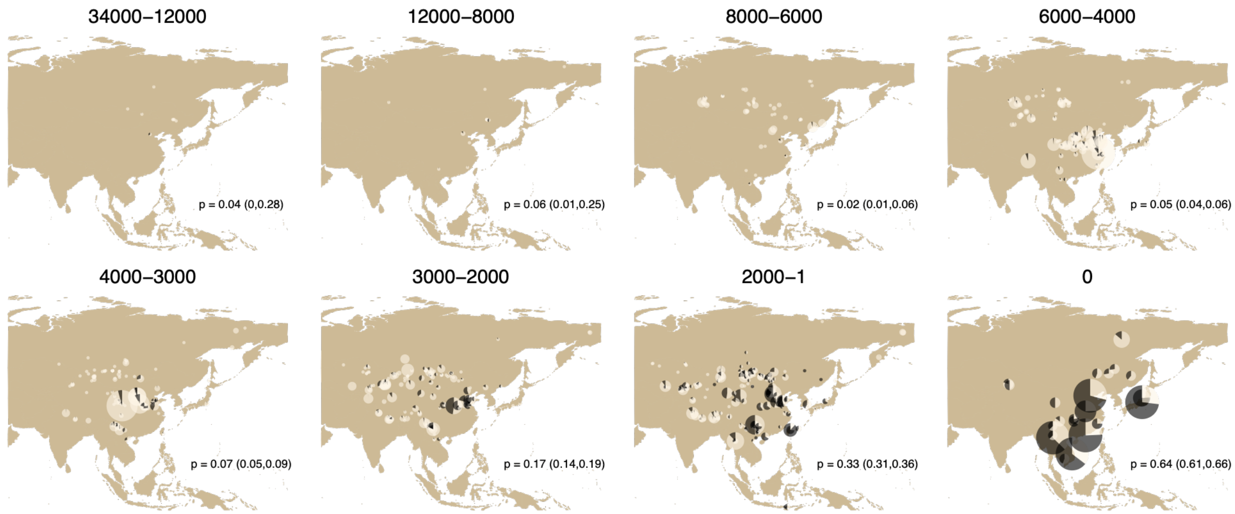


**Figure 14.** Derived allele frequency map based on imputed genotype dosages for rs1229984 (*ADH1B*).

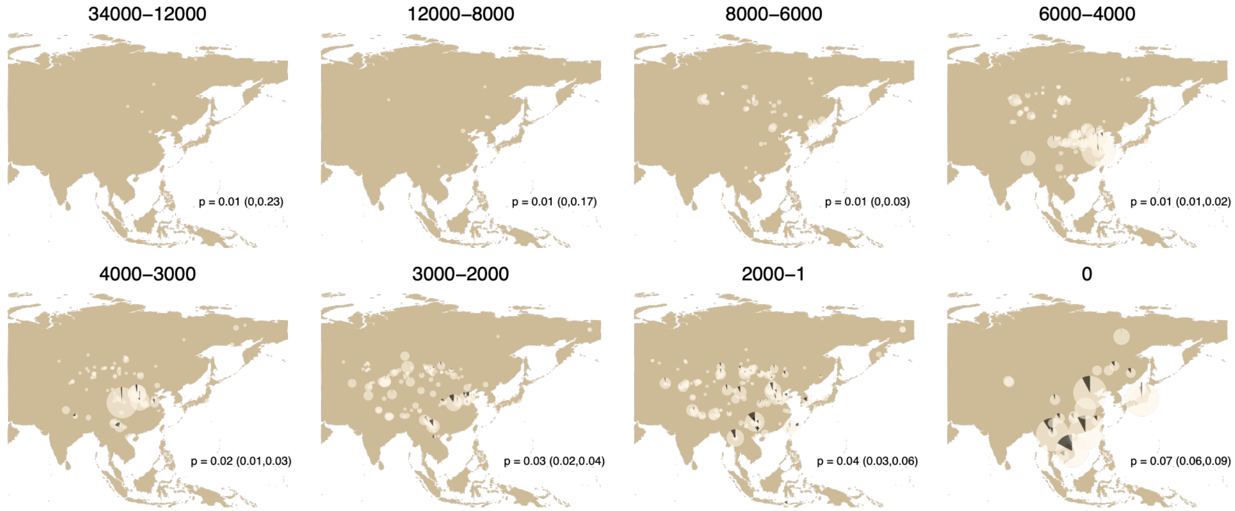


**Figure 15.** Derived allele frequency map based on imputed genotype dosages for rs671 (*ALDH2*).

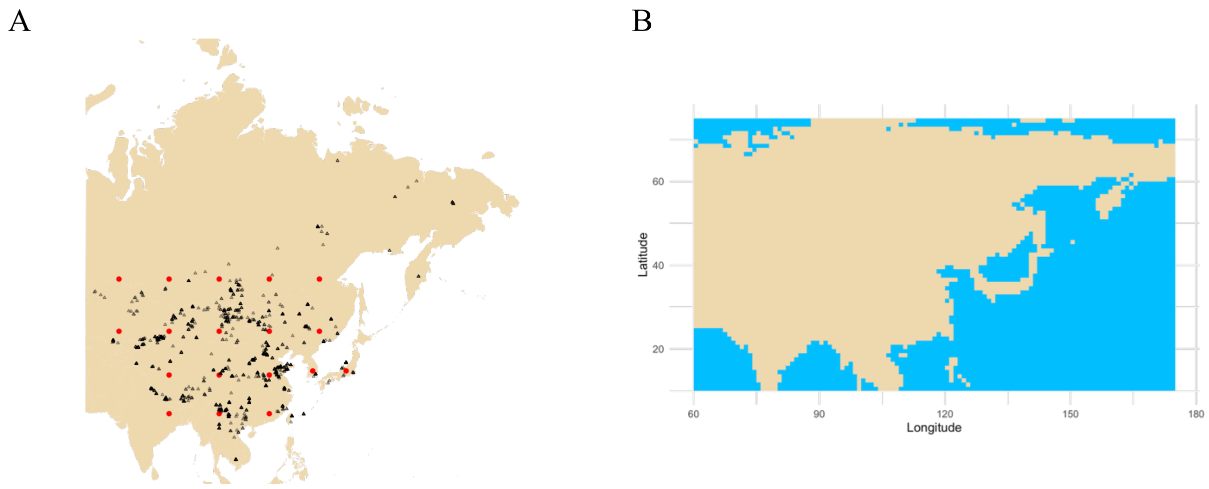


**Figure 16.** Spatiotemporal allele frequency reconstruction software configurations for (A) locations of potential points of allele origin represented by red dots (triangles represent locations of all individuals in the dataset) and (B) the topology of the analyzed landmass that was manipulated to create artificial land bridges connecting Japan and Taiwan to mainland East Asia.

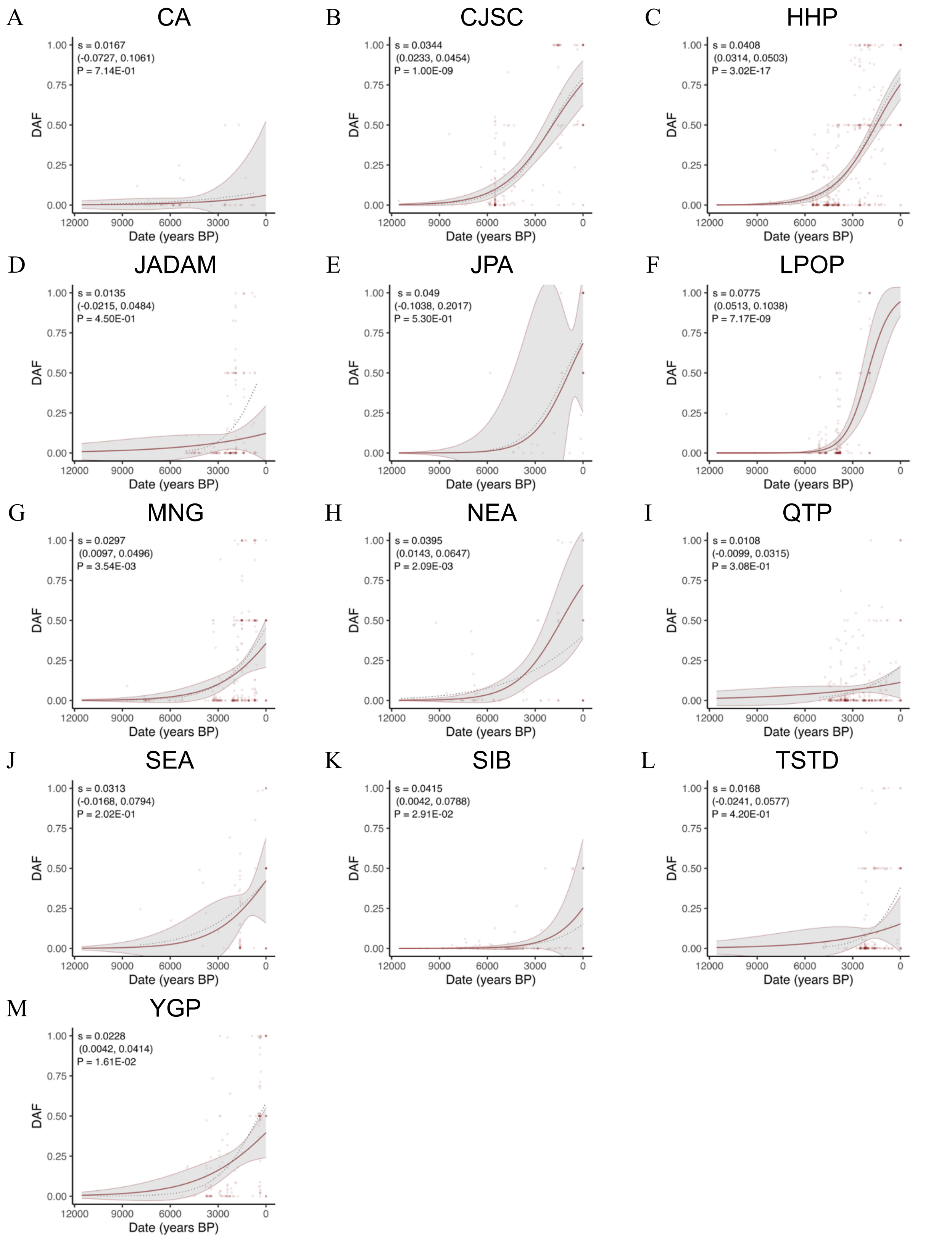


**Figure 17.** Allele frequency trajectory reconstructions for 48His allele of rs1229984 in ADH1B for (A) Central Asia, (B) Chang Jiang & Southern Coastal China & Sichuan Basin, (C) Huang

He Plain, (D) Junggar & Alashan Semi-deserts & Altai Mt., (E) Japanese Archipelago, (F) Loess Plateau & Ordos Plateau, (G) Mongolian Grasslands, (H) Northeast Asia, (I) Qinghai-Tibetan Plateau, (J) Southeast Asia, (K) Siberia, (L) Tianshan & Taklimakan Desert, and (M) Yunnan-Guizhou Plateau. The filled line with 95% CI envelope represents the regression with covariates including genomic PCs; the dashed line represents the prediction from a model with just date as predictor.

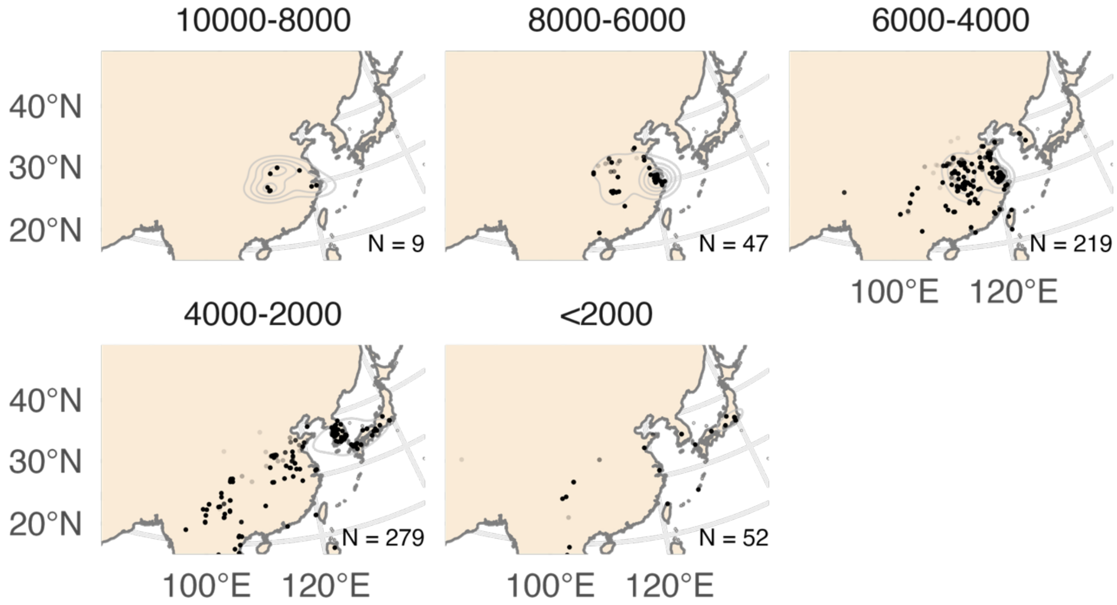


**Figure 18.** Kernel density estimates for rice farming intensity in East Asia during Holocene. Here and for the following plots, we based the estimates on relative contributions to the total biomass per site.

A

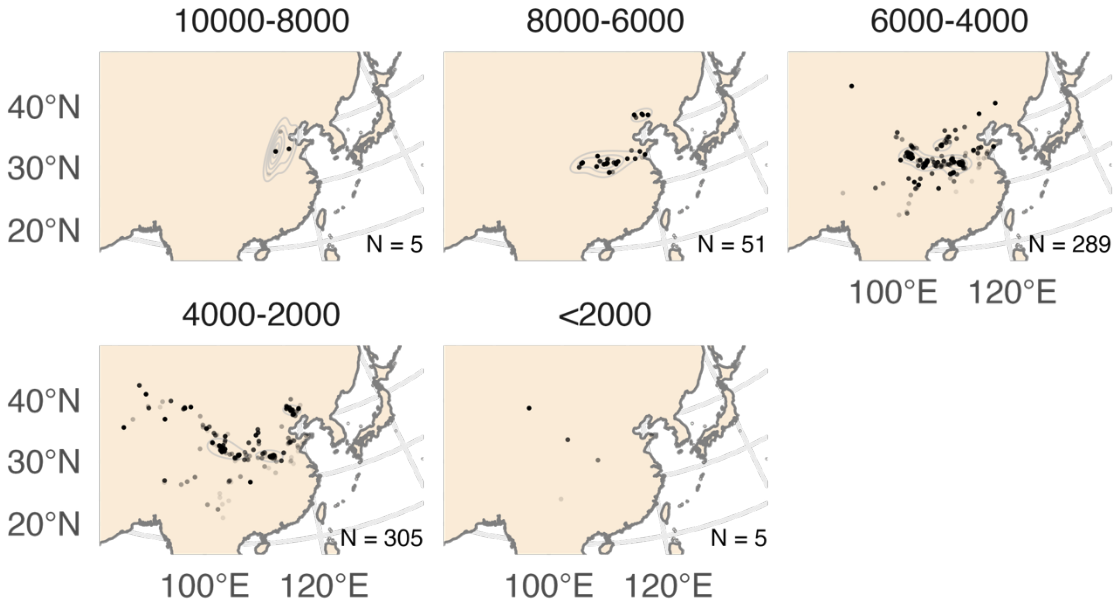

B

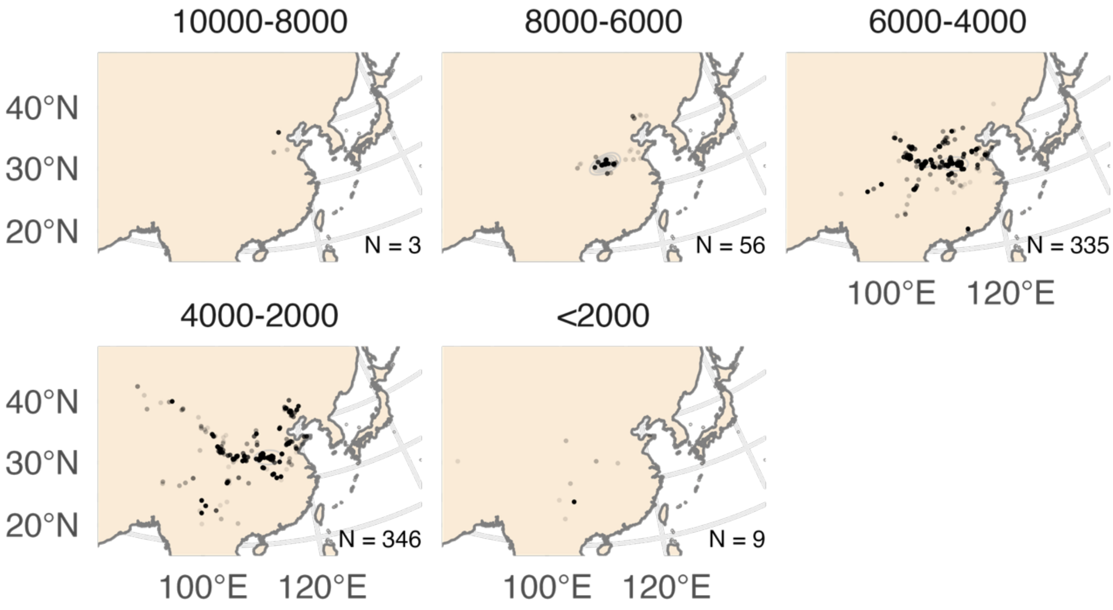


**Figure 19.** Kernel density estimates for (A) broomcorn and (B) foxtail millet sites.

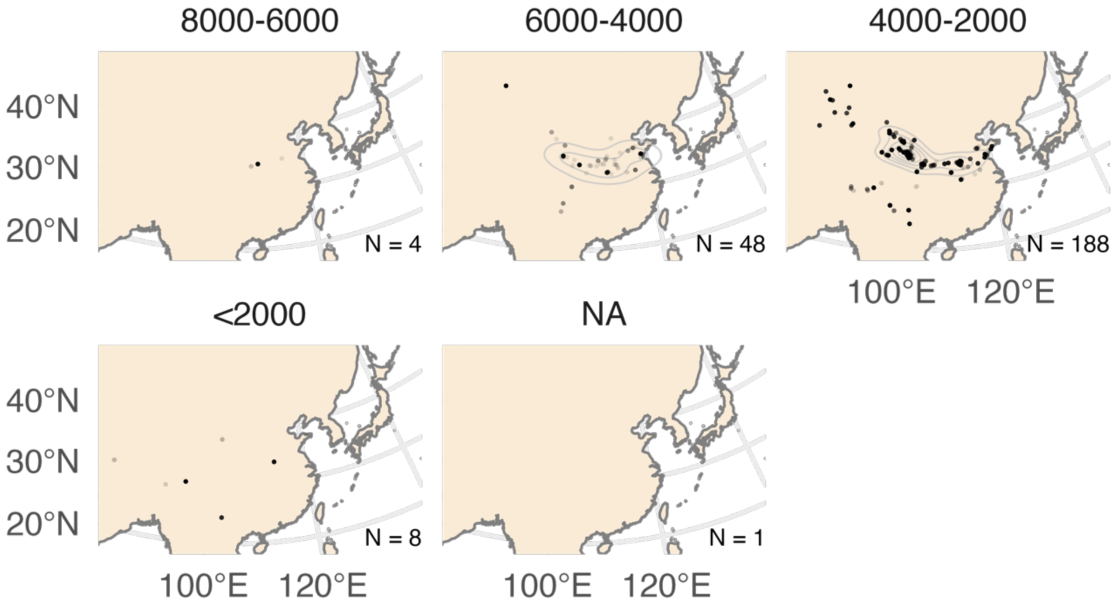


**Figure 20.** Kernel density estimates for wheat farming intensity in East Asia during Holocene.

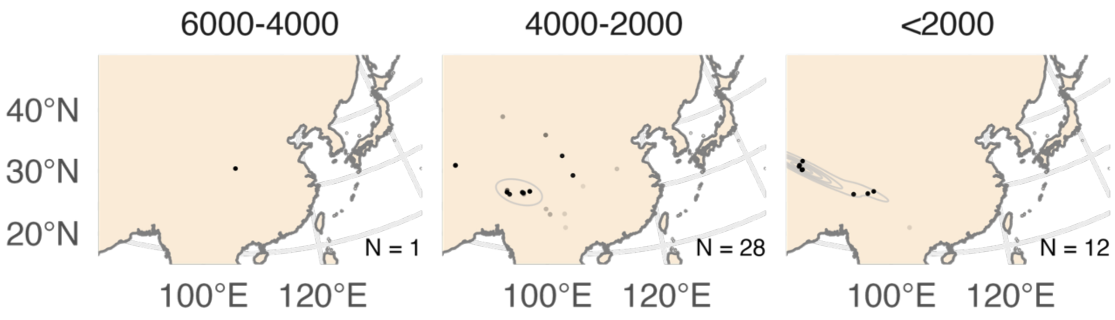


**Figure 21.** Kernel density estimates for barley farming intensity in East Asia during Holocene.

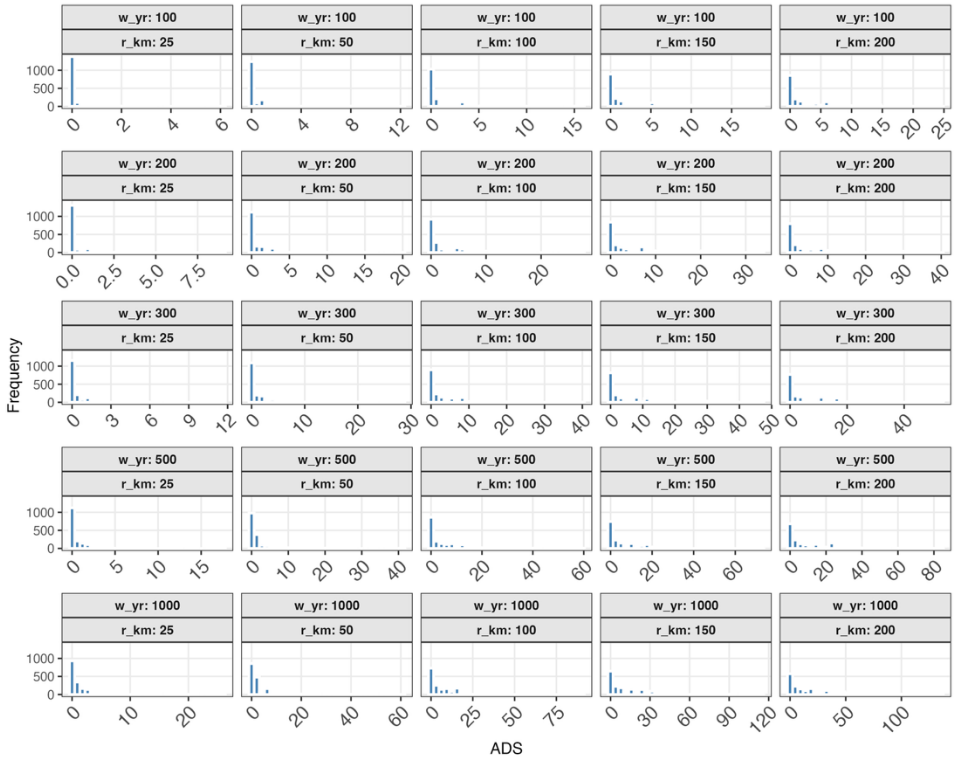


**Figure 22.** Distributions of archaeobotanical density scores of ancient individuals for different bandwidth parameter combinations.

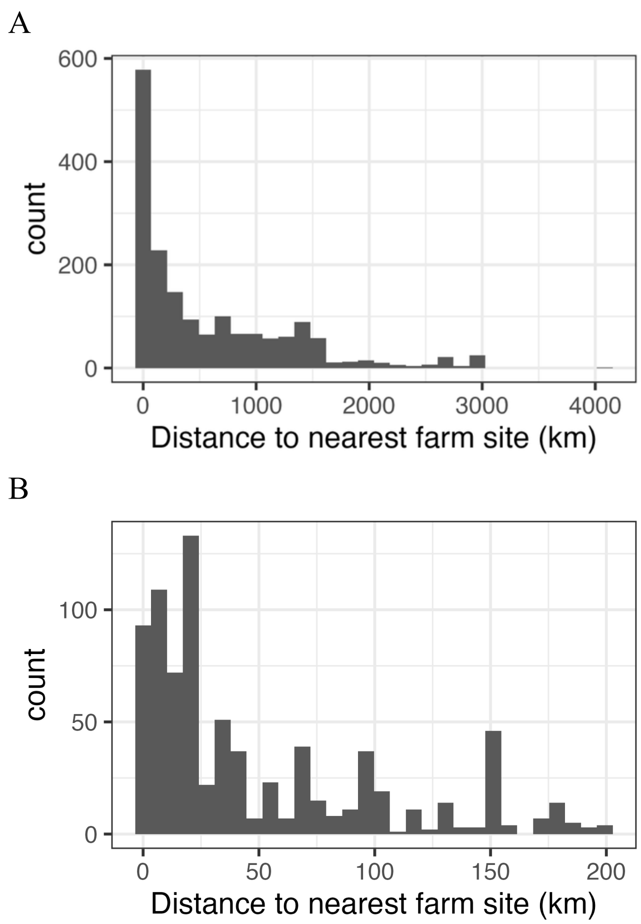


**Figure 23.** Histogram of the distances of the nearest archaeobotanical site to ancient individual that are dated to within +/- 1000 years of the individual's date for (A) all individuals and (B) individuals who are located at most 200 km away from the nearest site.

A

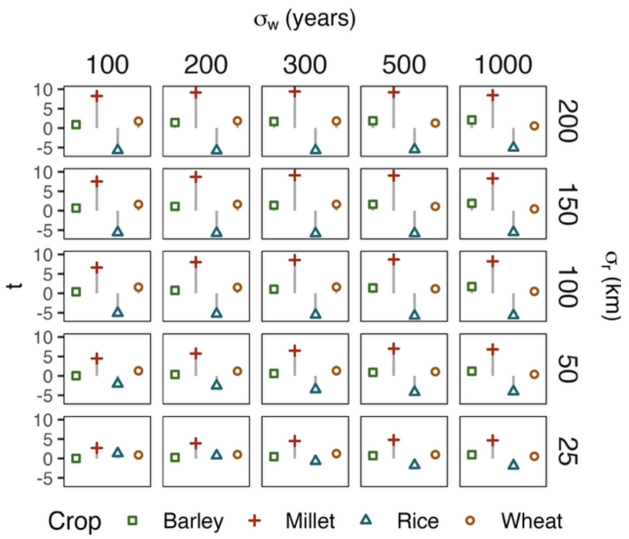


B

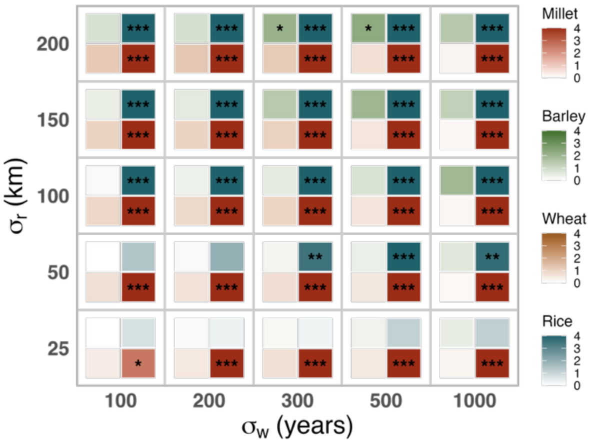


**Figure 24.** Crop specific ADS associations with rs174546 in *FADS2* and (B) heatmap of permutation-based P-values

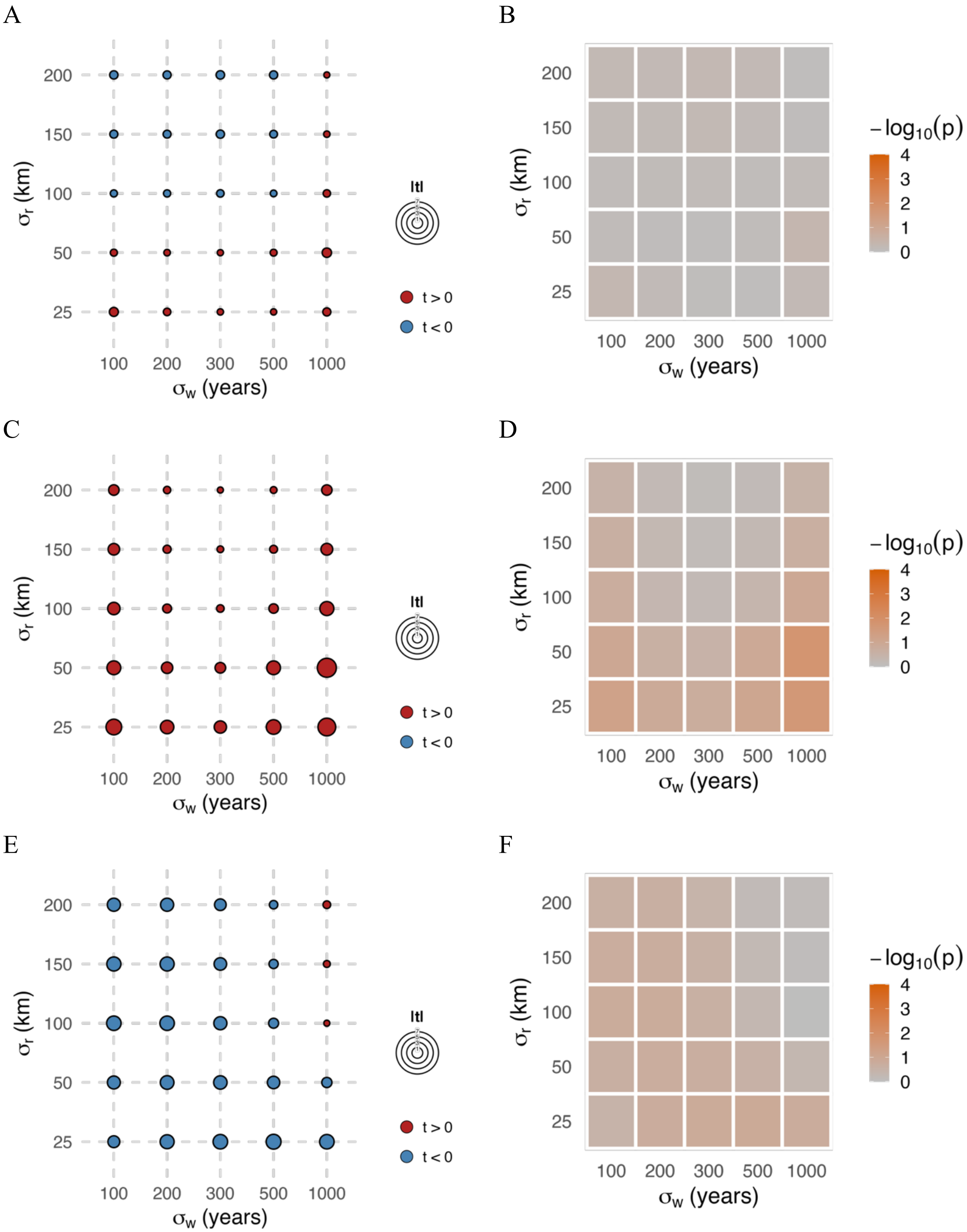
A
B
C
D
E
F
200
150
100
50
25
$\sigma_r$ (km)
100 200 300 500 1000
$\sigma_w$ (years)
|t|
t > 0
t < 0
$-\log_{10}(p)$
4
3
2
1
0

G

H

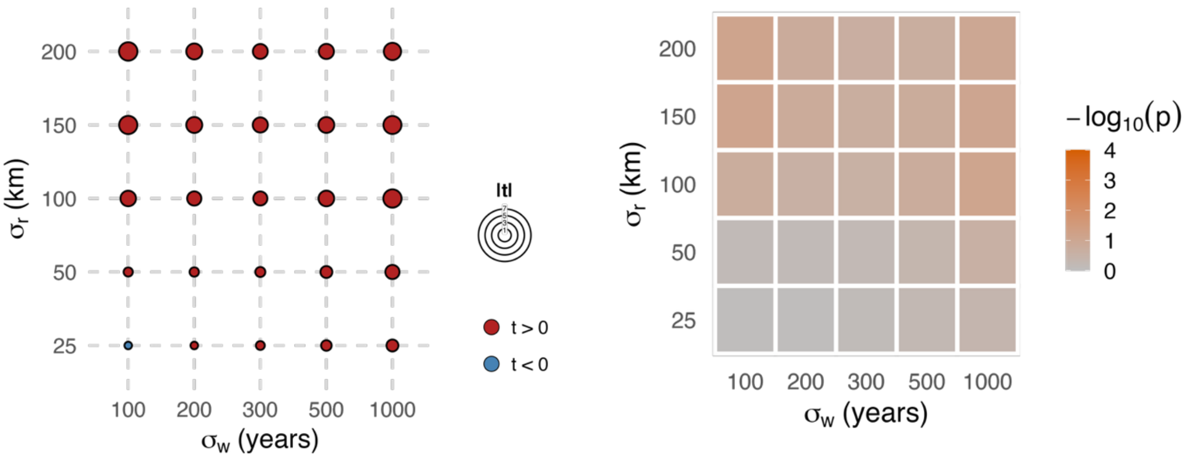


I J

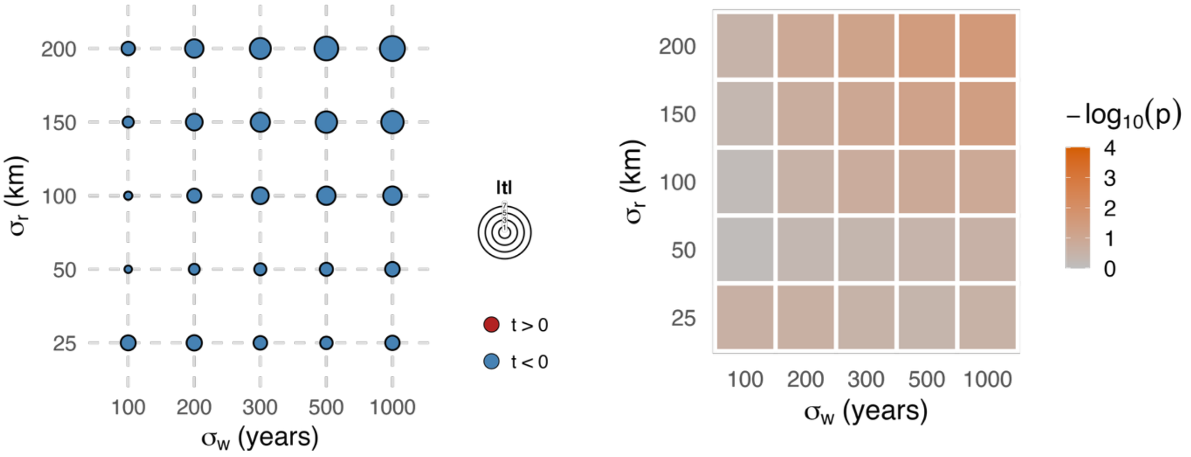


**Figure 25.** ADS association test statistics and permutation P-value heat map for (A, B) rs671 (*ALDH2*), (C, D) rs9258176 (*HLA-DQA2*), (E, F) rs79230913 (*OAS1*), (G, H) rs784133 (*RSPH4A*), and (I, J) rs6766450 (*STAG1*).

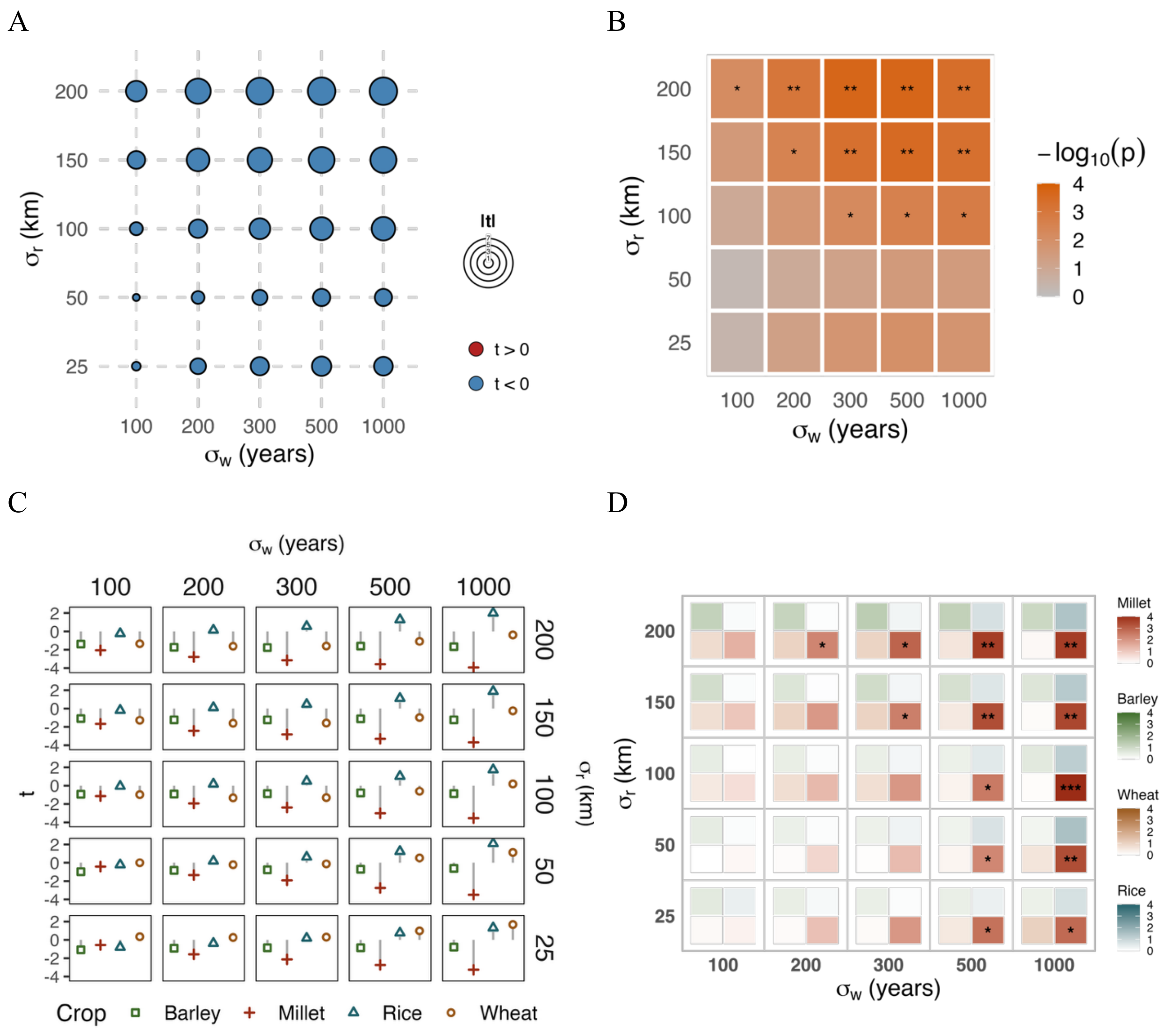


**Figure 26.** (A) ADS association test statistics for rs1800414 (*OCA2*) and (B) heatmap of permutation-based P-values. (C) Crop-specific ADS association test statistics and (D) heatmap of permutation-based P-values for each crop.

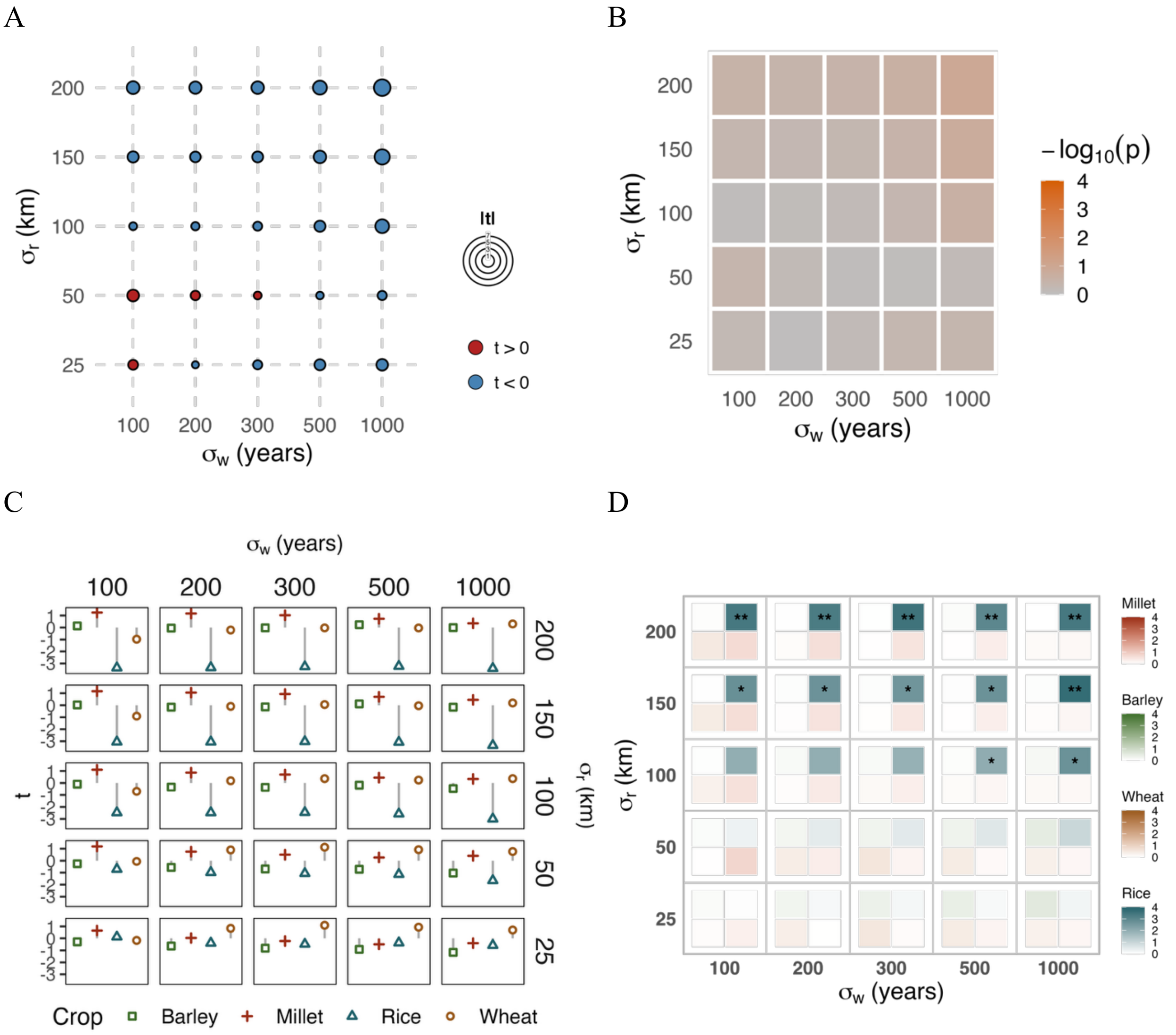


Figure 27. (A) ADS association test statistics for rs1047781 (*FUT2*) and (B) heatmap of permutation-based P-values. (C) Crop-specific ADS association test statistics and (D) heatmap of permutation-based P-values for each crop.

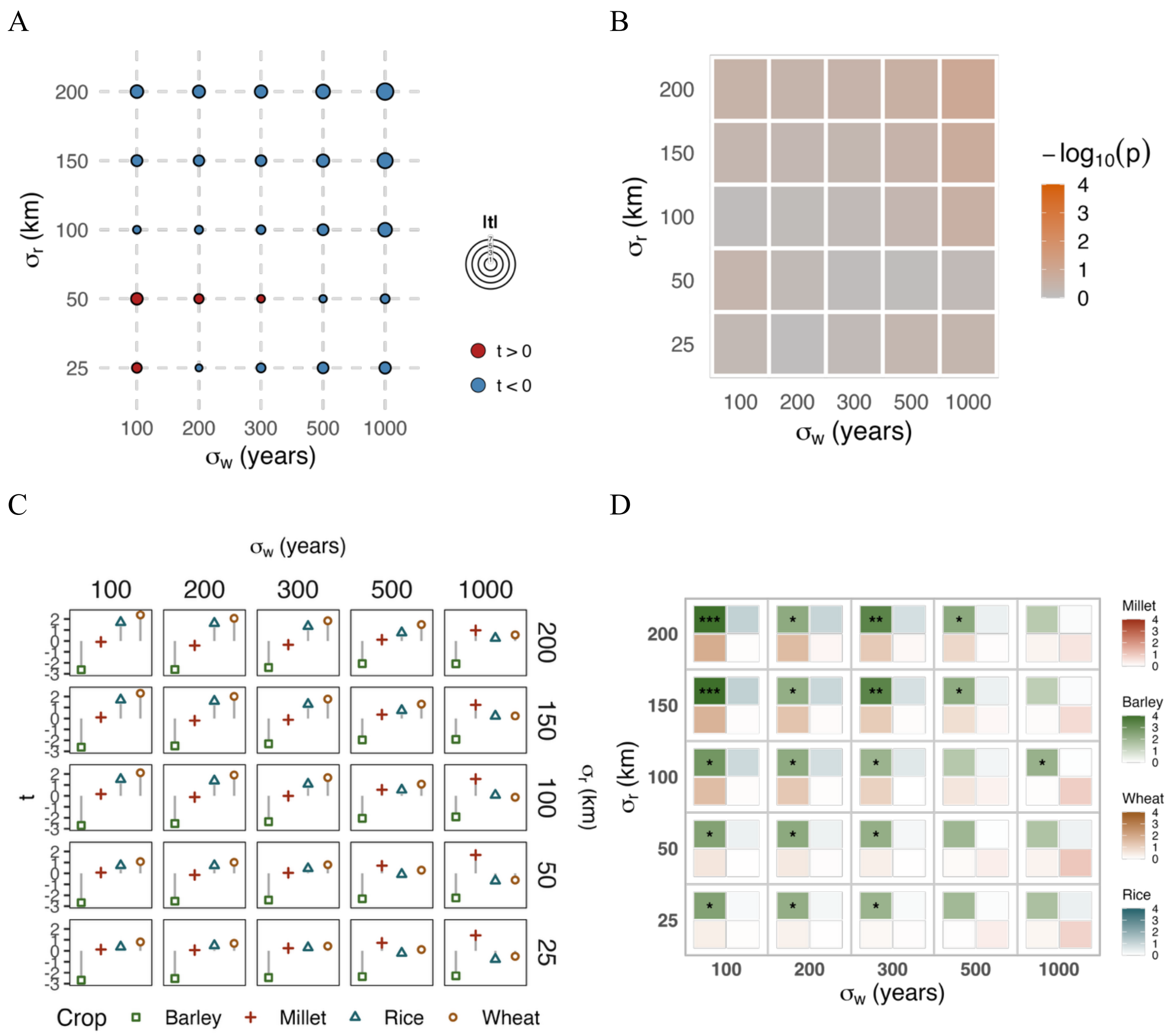


**Figure 28.** (A) ADS association test statistics rs784133 (*RSPH4A*) and (B) heatmap of permutation-based P-values. (C) Crop-specific ADS association test statistics and (D) heatmap of permutation-based P-values for each crop.

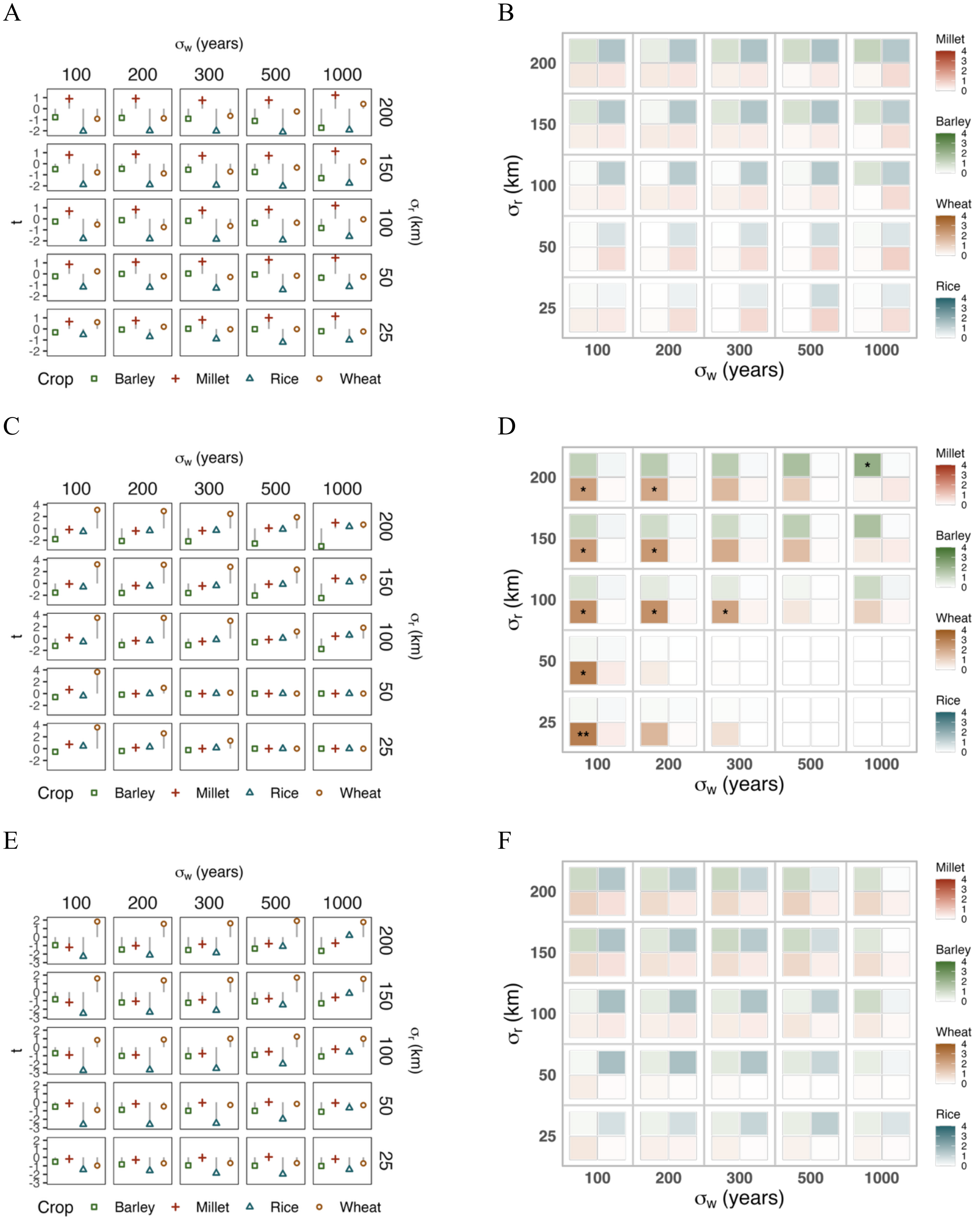
A
σw (years)
100 200 300 500 1000
t
σr (km)
200 150 100 50 25
Crop Barley Millet Rice Wheat
B
σr (km)
σw (years)
Millet
Barley
Wheat
Rice
C
D
E
F

G

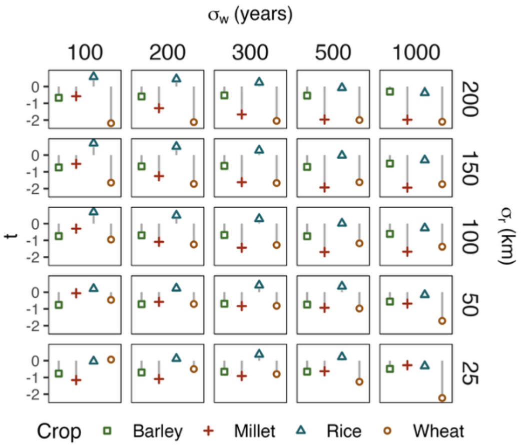


H

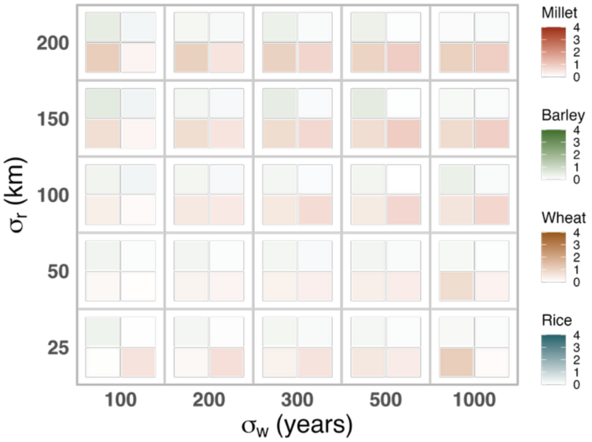


**Figure 29.** Crop-specific ADS association test statistics and permutation P-value heat map for (A, B) rs671 (*ALDH2*), (C, D) rs9258176 (*HLA-DQA2*), (E, F) rs79230913 (*OAS1*), and (G, H) rs6766450 (*STAG1*).

**Supplementary Note 1: Evaluation of imputation accuracy in ancient East Asian genomes**

While approximately one third of the 1000 Genomes Project reference panel was of East Asian origin, the particular ancestry groups represented in 1000 Genomes might not be fully representative of the diversity in East Asia that was present in the past. In addition, the available ancient nuclear data from East Asia are generally of lower coverage compared to data from Europe, and previous studies that have quantified the performance of imputation of ancient genomes indicate issues with bias in the imputed genotypes particularly for lower coverage samples[1,2]. Specifically, imputation on low coverage individuals can induce them to resemble the average of the reference panel. To investigate the genotype accuracy of imputation with GLIMPSE2, we performed downsampling experiments with higher coverage capture sample data that were generated from our lab. Our results examining higher coverage capture samples from East Asia were qualitatively consistent with Mota et al.'s in terms of their imputation accuracy for ancient West Eurasian and Siberian individuals. We only observed relatively high genotype accuracy ($r^2 >$ 0.9) for imputation of common variants in East Eurasian capture genomes of at least 1x or 2x coverage depending on the individual (Fig. 30). However, for 0.5x coverage or even 0.2x coverage we generally saw that $r^2 > 0.5$. The diminished genotype accuracy did result in bias for downstream analyses of relatedness, such as in PCA. The lowest coverage versions of the original samples displayed the furthest distance on PCA from the unimputed version, and indeed, there was a general trend of the imputed genome moving towards the average genotype, at the center of PC space, with decreasing coverage (Fig. 31). We also investigated imputation performance between the high coverage 1000 Genomes reference panel and the harmonized 1000 Genomes + HGDP panel[3] as part of a preliminary study. In general, we observed only a marginal improvement in imputation accuracy with the larger and more diverse panel at most increasing the $r^2$ by less than 0.03 and the biggest improvements being in the MAF range of 0.02 to 0.10 for most coverages (Fig. 32). Furthermore, the 1000 Genomes + HGDP panel filtered out some well-established adaptive variants including rs1229984, so we decided to use the 1000 Genomes reference panel for this reason and the minimal gains in accuracy for the larger panel.

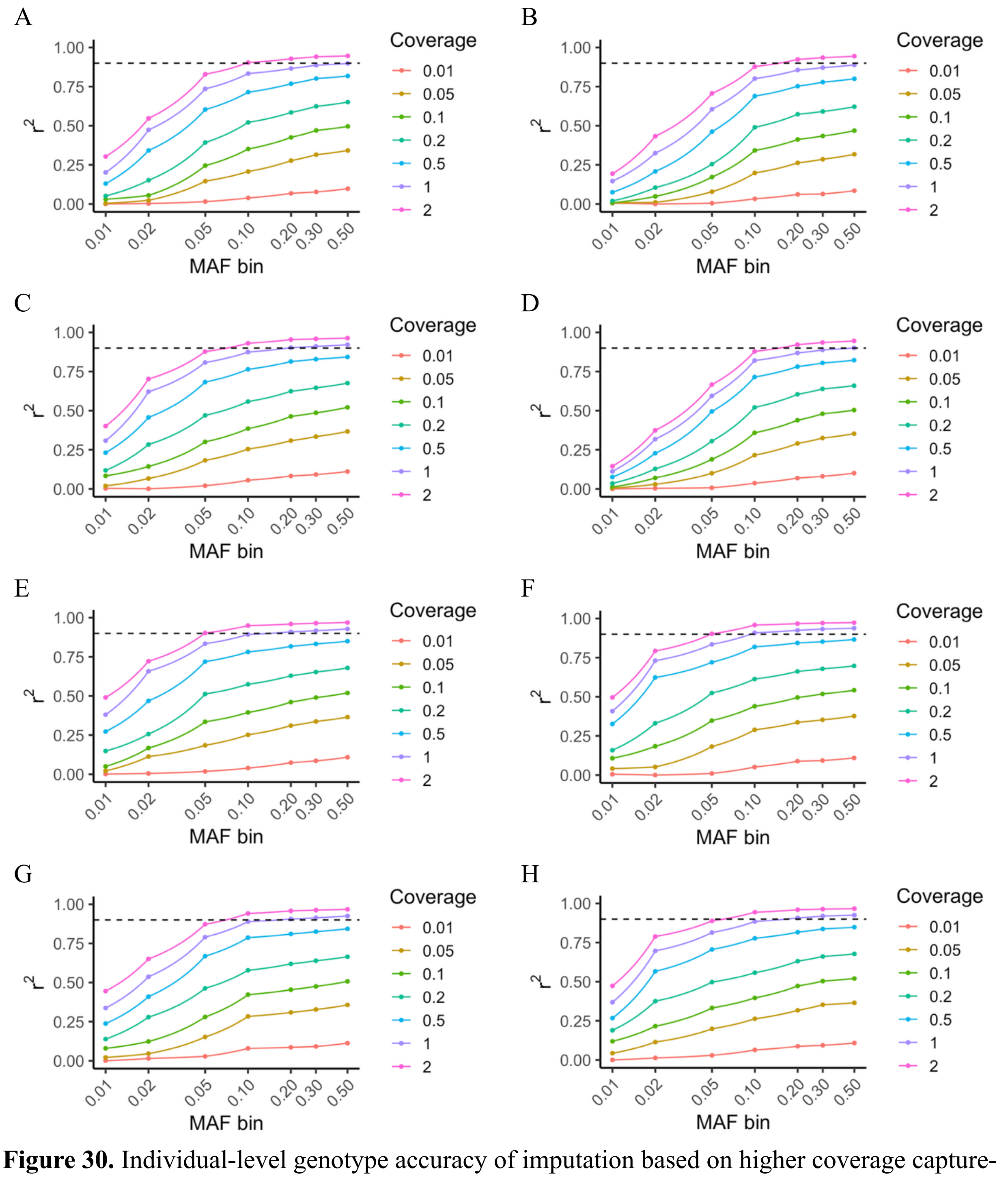


**Figure 30.** Individual-level genotype accuracy of imputation based on higher coverage capture-enriched ancient genomes from East Asia: (A) AR33k, (B) Xiaogao, (C) Nagqu1.4k, (D) DHL_M2, (E) Gebusailu2.3k, (F) Wuzhuangguoliang, (G) Sangdalongguo, and (H) Shigatse2.1k. The dotted line indicates $r^2 = 0.9$. Each tick for the MAF bin indicates SNPs with an MAF greater than the previous tick and up to the labelled value (e.g., the 0.50 bin represents SNPs with MAF > 0.30 and ≤ 0.50).

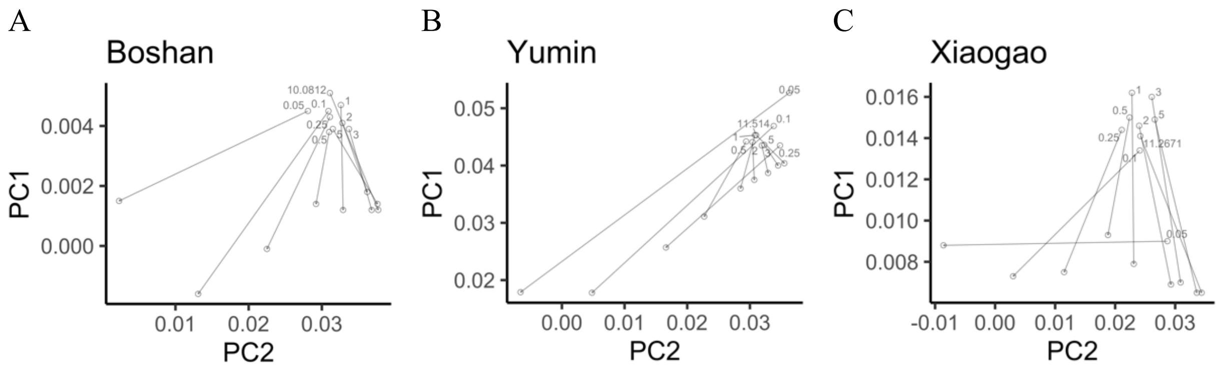


**Figure 31.** Comparisons of the PCA positions (projected onto PCs generated from East Asian HGDP individuals) of original and downsampled data for three relatively high coverage ancient genomes with the arrows pointing from unimputed, for which standard haploid genotype calling was performed, to imputed data. Labels indicate the coverage of from the original samples to the downsampled amounts (5x, 3x, 2x, 1x, 0.5x, 0.25x, 0.1x, 0.05x).

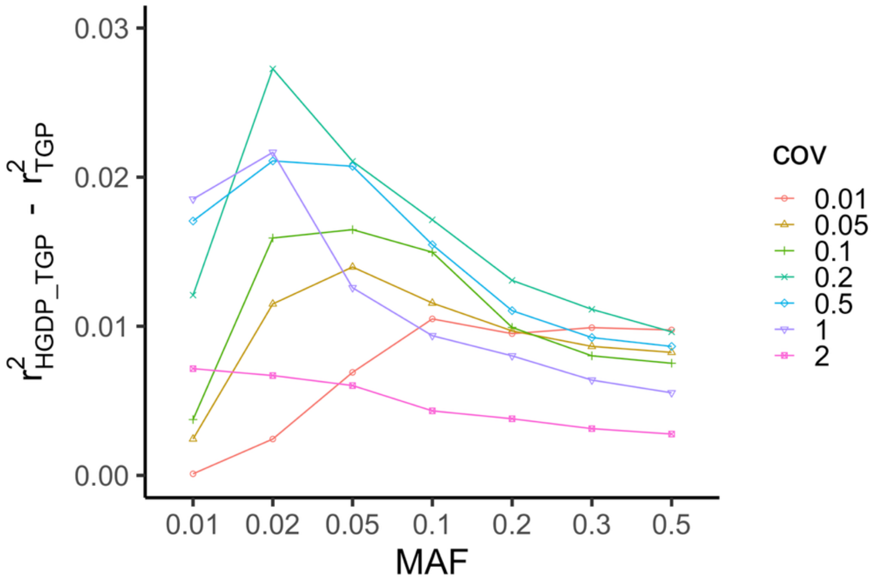


**Figure 32.** Evaluation of imputation accuracy comparing the high coverage 1000 Genomes reference panel to the jointly called 1000 Genomes and HGDP reference panel.
